# Mono- to tetra-alkyl ether cardiolipins in a mesophilic, sulfate-reducing bacterium identified by UHPLC-HRMS^n^: A novel class of membrane lipids

**DOI:** 10.1101/2024.03.20.586015

**Authors:** Ellen C. Hopmans, Vincent Grossi, Diana X. Sahonero Canavesi, Nicole J. Bale, Cristiana Cravo-Laureau, Jaap S. Sinninghe Damsté

## Abstract

The composition of membrane lipids varies in a number of ways as adjustment to growth conditions. Variations in head group composition and carbon skeleton and degree of unsaturation of glycerol-bound acyl or alkyl groups results in a high structural complexity of the lipidome of bacterial cells. We studied the lipidome of the mesophilic, sulfate-reducing bacterium, *Desulfatibacillum alkenivorans* strain PF2803^T^ by ultra-high-pressure liquid chromatography coupled with high-resolution tandem mass spectrometry (UHPLC-HRMS^n^). This anaerobic and hydrocarbon-utilizing bacterium has been previously shown to produce high amounts of mono- and di-alkyl glycerol ethers as core membrane. Our analyses revealed that these core lipids occur with phosphatidylethanomamine (PE) and phosphatidylglycerol (PG) head groups, representing each approximately one third of the phospholipids. The third class was a novel group of phospholipids, i.e. cardiolipins (CDLs) containing one (monoether/triester) to four (tetraether) ether-linked saturated straight-chain or methyl-branched alkyl chains. Tetraether CDLs have been shown to occur in archaea (with isoprenoid alkyl chains) but have not been previously reported in the bacterial Domain. Structurally related CDLs with one or two alkyl/acyl chains missing, so-called monolyso- and dilyso-CDLs, were also observed. The potential biosynthetic pathway of these novel CDLs was investigated by examining the genome of *D. alkenivorans*. Three CDL synthases were identified; one catalyzes the condensation of two PGs, the other two are probably involved in the condensation of a PE with a PG. A heterologous gene expression experiment showed the *in vivo* production of dialkylglycerols upon anaerobic expression of the glycerol ester reductase enzyme of *D. alkenivorans* in *E. coli*. Reduction of the ester bonds probably occurs first at the *sn*-1 and subsequently at the *sn*-2 position after the formation of PEs and PGs, since PGs possess a much higher percentage of ether bonds than PEs.

## INTRODUCTION

Bacterial and eukaryotic cellular membranes are composed of a lipid bilayer made of intact polar lipids, commonly diacyl glycerols with a polar head group such as phosphatidylglycerol (PG), phosphatidylserine (PS) or phosphatidylethanomamine (PE). Cardiolipins (CDLs), formally 1,3-bis(sn-3’-phosphatidyl)-*sn*-glycerols also called bisphosphatidylglycerols, are a particular class of phospholipids with a somewhat unusual structure. They can be considered as dimeric phospholipids in which two 1,2-diacyl-*sn*-glycero-3-phosphoryl moieties are bound through a third glycerol moiety (see Figure 1 for structures), resulting in phospholipids with four instead of two acyl chains with varying chain length and degree of unsaturation, resulting in complex mixtures (Oemer et al., 2018). CDLs are known to be involved in the structural organization of membranes, protein interactions, enzyme functioning, and osmoregulation (Schlame, 2008). Their biosynthesis in bacteria is encoded by cardiolipin synthases (Cls), enzymes belonging to the phospholipase D (PLD) superfamily or to the CDP-alcohol phosphotransferase superfamily (Sandoval-Calderon et al., 2009). CDLs occur widely in membranes of eukaryotes and bacteria but have also been reported in halo(alkali)philic and methane-metabolizing archaea, where they occur as unusual CDLs with four ether-bound isoprenoidal alkyl chains (i.e., so-called tetraethers) instead of the common esterified acyl chains found in bacterial and eukaryotic phospholipids (Lattanzio et al, 2002; Corcelli, 2009; Angelini et al., 2012; Yoshinaga et al., 2012; Bale et al., 2019). CDLs with core lipids containing plasmalogens, i.e. 1-O-alk-1’-enyl, 2-acyl glycerolipids, have been reported in anaerobic bacteria and myxobacteria (see Goldfine, 2017 for a review) but these CDLs contain two ester bonds and, in addition, two vinyl ether moieties.

**Figure 1.**
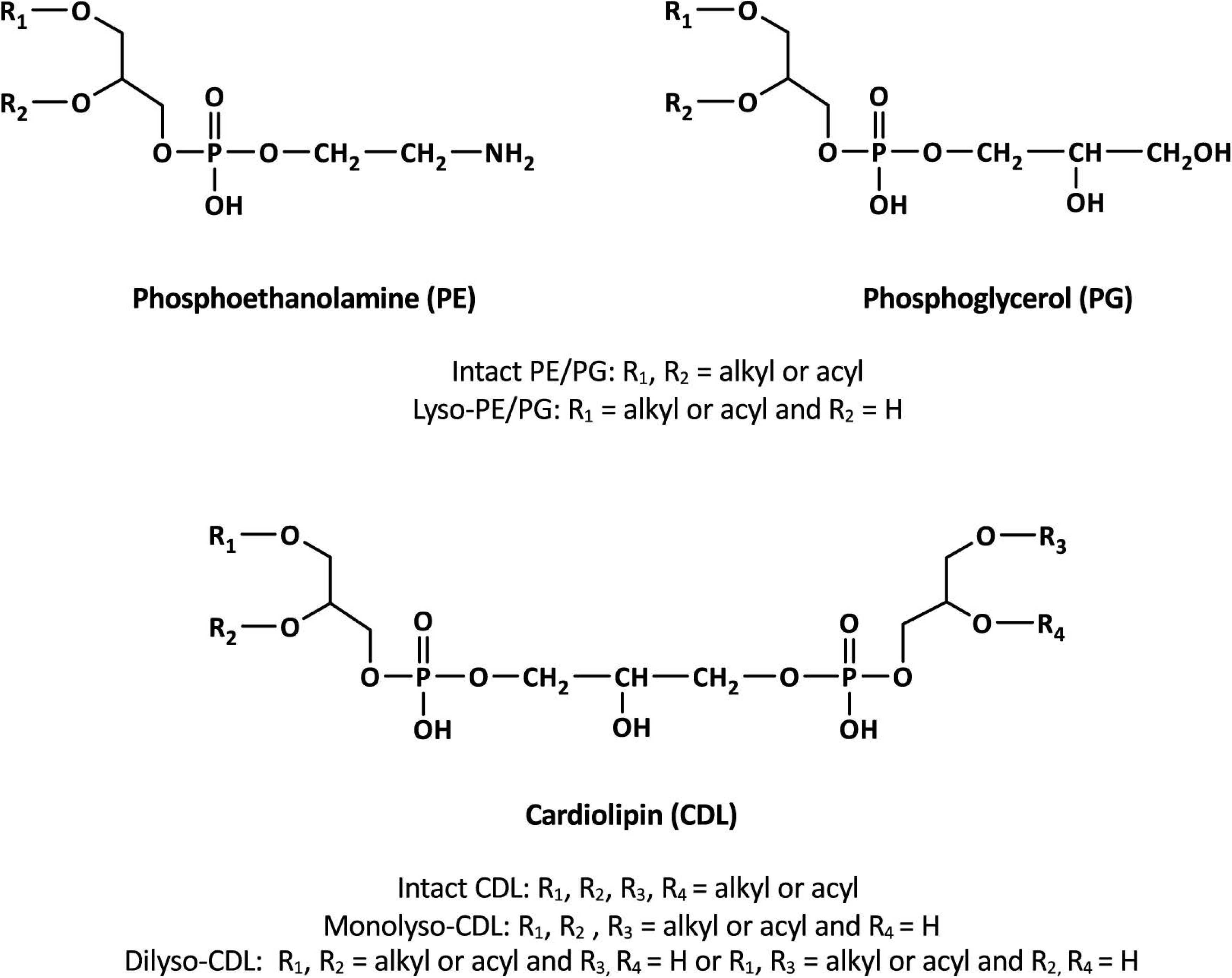
General structures of membrane phospholipids discussed in the text.

Although saturated ether lipids are typically considered to be of unique archaeal origin, some mesophilic and thermophilic bacteria have the capacity to biosynthesize monoalkyl/monoacyl- and dialkyl-glycerol ether lipids (e.g., Langworthy et al., 1983; Huber et al., 1996; Sturt et al., 2004; Ring et al., 2006; Grossi et al., 2015; Vinçon-Laugier et al., 2016; Sinninghe Damsté et al., 2004, 2007, 2014, 2023). Such non-isoprenoid alkylglycerols are increasingly being recognized in bacterial isolates and are widespread in the environment (see Grossi et al., 2015 for review). The biosynthesis of alkylglycerols in bacteria has been described to be catalyzed either by the ether lipid biosynthesis (ElbD, Lorenzen et al., 2014) or glycerol ester reductase (Ger; Sahonero-Canavesi et al., 2022) proteins. Heterologous gene expression experiments showed the *in vivo* production of different monoalkylglycerols upon anaerobic expression of the Ger enzyme of the alkylglycerol-producing sulfate-reducer *Desulfatibacillum alkenivorans* in *E. coli*. It was demonstrated that Ger converts an *sn*-1 ester bond into an *sn*-1 ether bond and that the ether-bound alkyl chains are based on the major fatty acid components of the expression host (Sahonero Canavesi et al., 2022). However, although these findings could explain the occurrence of *sn-*1-alkyl ether bonds in a wide variety of bacteria, it did not provide evidence for the biosynthesis of glycerol diethers, such as those observed in *D. alkenivorans* (Grossi et al, 2015), suggesting that a second enzyme would be responsible for the formation of the alkyl ether bonds at the *sn*-2 position of the glycerol moiety.

Here, we report the composition of the lipidome of the mesophilic sulfate-reducing bacterium, *D. alkenivorans* strain PF2803^T^. Previous analysis of the membrane core lipids (obtained after hydrolysis of the phospholipids) of this bacterium and a phylogenetically closely related species (*D. aliphaticivorans* strains CV2803^T^) showed that their membrane lipids deviate from those of most other mesophilic bacteria because they contain high amounts of mono- and di-alkyl glycerol ethers (Grossi et al., 2015; Vinçon-Laugier et al., 2016; 2017). However, their phospholipid composition remained unknown. We applied UHPLC-HRMS^n^ to characterize the lipidome of *D. alkenivorans* strain PF2803^T^ in detail and came across a class of bacterial phospholipids that has not yet been characterized, i.e. CDLs containing one (monoether/triester) to four (tetraether) ether-linked saturated alkyl chains, resulting in an even more complex distribution of bacterial CDLs as previously reported (Oemer et al., 2018). Corresponding CDLs with one or two alkyl/acyl chains missing, so-called monolyso- and dilyso-CDLs, were also observed. The potential biosynthetic pathway of these novel CDLs was investigated by examining the genes encoding the proteins potentially involved.

## MATERIAL AND METHODS

### Bacterial Cultivation

The mesophilic sulfate-reducing bacterium *D. alkenivorans* strain PF2803^T^ was previously isolated from oil-polluted marine sediment as described by Cravo-Laureau et al. (2004). The strain belongs to the family *Desulfatibacillaceae* of the Thermodesulfobacteriota (Waite et al., 2020), formerly classified as a deltaproteobacterium. *D. alkenivorans* is a hydrocarbonoclastic bacterium capable of utilizing *n*-alk-1-enes in addition to more classical carbon substrates (Cravo-Laureau et al., 2004; Grossi et al., 2015; Vinçon-Laugier et al., 2016); the most efficient growth is generally observed with C_14_ to C_17_ substrates. Strain PF2803^T^ was grown under optimal growth conditions (30°C, pH 6.8, [NaCl] 10 g/L) in 100 ml of defined anoxic sulfate-reducing medium with *n*-hexadec-1-ene as the sole carbon and energy source. At the end of the exponential-growth phase, cells were harvested by filtration on 0.22 µm glass microfiber filters (GF/B; Whatman) and kept frozen before lipid analysis.

### Extraction and HPLC-HRMS^n^ Analysis of Phospholipids

Filtered cells were extracted using a modified Bligh and Dyer method using a mixture of methanol/dichloromethane/phosphate buffer (2:1:0.8, v/v/v), and extracts were then analyzed using a normal phase UHPLC-ESI/HRMS^n^ orbitrap method, as described in Bale et al. (2019). Phospholipid identification was based on exact mass (mass tolerance 3 ppm) and interpretation of MS^2^ fragmentation spectra. Relative abundances of phospholipids were determined by peak integration of mass chromatograms of the [M+H]^+^, [M+H-H_2_O)]^+^, [M+NH]^+^ [M+Na]^+^ ions as appropriate, and of the ^13^C isotope peak for the [M+H]^+^ ions in the case of the CDLs. It is important to note that different phospholipid classes exert a different MS response, and the MS response can even differ within a phospholipid class depending on the chemical structure of the core lipid (chain length, methyl branching, ether vs. ester bond). Due the analytical method used, the relative abundance of PGs and CDLs are likely underestimated due to their relatively poor ionization and loss of the molecular signal due to in-source fragmentations. The phospholipid distributions obtained do not take these differences into account and are based on the relative abundance of the lipids as measured.

### Genomic Methods and Phylogenetic Tree Construction

The genes encoding for the proteins involved in ether formation and cardiolipin biosynthesis were obtained from the UniProt database (The UniProt Consortium, 2023) or identified by protein Blast Searches (pBLAST). For the ether biosynthetic genes, a protein homologue to the confirmed Ger enzyme of the diether-producing *D. alkenivorans* (SHJ90043.1) were used as query. Similarly, the Cls proteins for the cardiolipin biosynthesis were identified with pBLAST using the ClsA/ClsB/ClsC (P0A6H8/P0AA84/P75919) from *E. coli* as query. For the multiple sequence amino acid analysis, the sequences were retrieved from the literature or the Uniprot database (The UniProt Consortium, 2023). For phylogenetic tree construction the amino acid sequences of all the characterized bacterial, archaeal and the proposed eukaryotic proteins were aligned with MAFFT, with a gap extension penalty of 0.123 and a gap open penalty of 1.53. This alignment was used as input for constructing the phylogenetic tree, using the PhyML in the online platform NGPhylogeny.fr (Lemoine et al., 2019). The phylogenetic tree was visualized with iTOL (Letunic and Bork, 2021). To examine the enzymatic activity of the additional glycerol ether reductase enzymes, the gene coding for a Ger homologue (SHK01260.1) was commercially synthesized (Eurofins, Germany), sub cloned in pCDFDuet-1 and expressed in *E. coli* strain BL21 DE3 aerobically and anaerobically, using the methods as described in Sahonero-Canavesi et al. (2022).

## RESULTS

### Dialkyl, Monoalkyl/Monoacyl, and Diacyl Composition of phospholipids

Figure 2 shows a partial base peak chromatogram (BPC) of the Bligh-Dyer extract of *D. alkenivorans* containing the phospholipids. The BPC is dominated by overlapping distributions of phospholipids with phosphoethanolamine (PE) and phosphoglycerol (PG) head groups and dialkylglycerol (DEG), monoalkyl/monoacylglycerol (AEG) and diacylglycerol (DAG) core lipids. PGs were detected in slightly higher proportions than PEs (Figure 3A). Dialkylglycerides with a PE or PG head group (DEG-PG and -PE) eluted between 19 and 21 min, alkyl/acylglycerides with a PE or PG head group (AEG-PG and -PE) eluted between 19.5 and 21.5 min, and diacylglycerides with PE- and PG head group (DAG-PG and -PE) eluted between 20 and 22.5 min. In addition, monoalkyl- and monoacyl-glycerides (MEG and MAG, respectively) with a PE or PG head group were detected. These lyso-lipids elute later, between 26 and 30 min.

**Figure 2.**
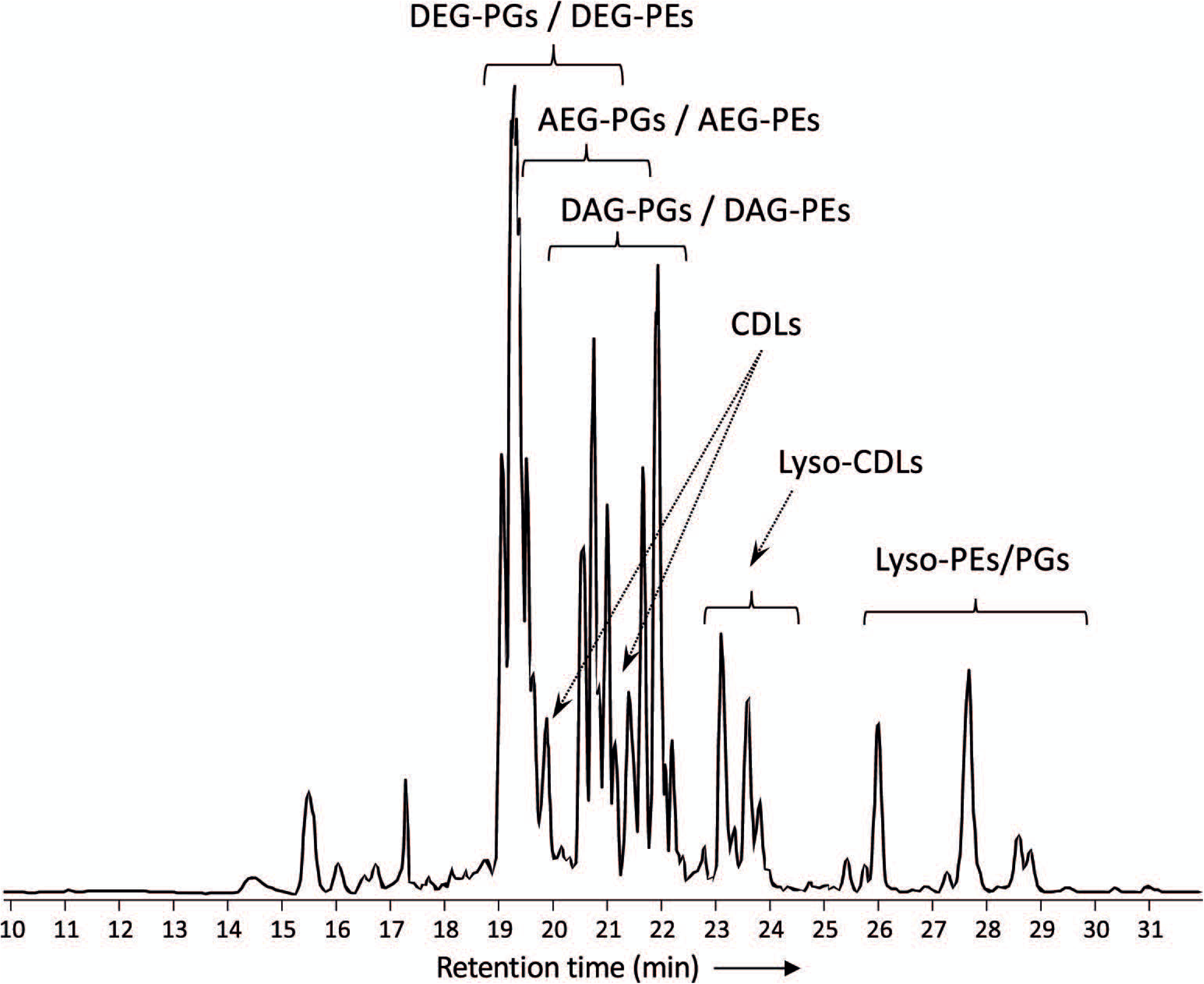
Partial base peak chromatogram (*m/z* 375-2000) of the total lipid extract of *D. alkenivorans* strain PF2803T grown on C_16_ *n*-alk-1-ene. Indicated are clusters of overlapping phospholipids (DEG = dialkylglyceride, AEG = monoalkyl/monoacylglyceride, DAG = diacylglyceride, PG = phosphoglycerol, PE = phosphoetanolamine) and high molecular weight compounds identified as alkyl ether cardiolipins (CDLs). Lyso-PEs/PGs = mixture of mono- alkyglyceride (MEG) and monoacylglyceride (MAG) PEs/PGs. Lyso-CDLs = mixture of mono- and dilyso-cardiolipins.

**Figure 3.**
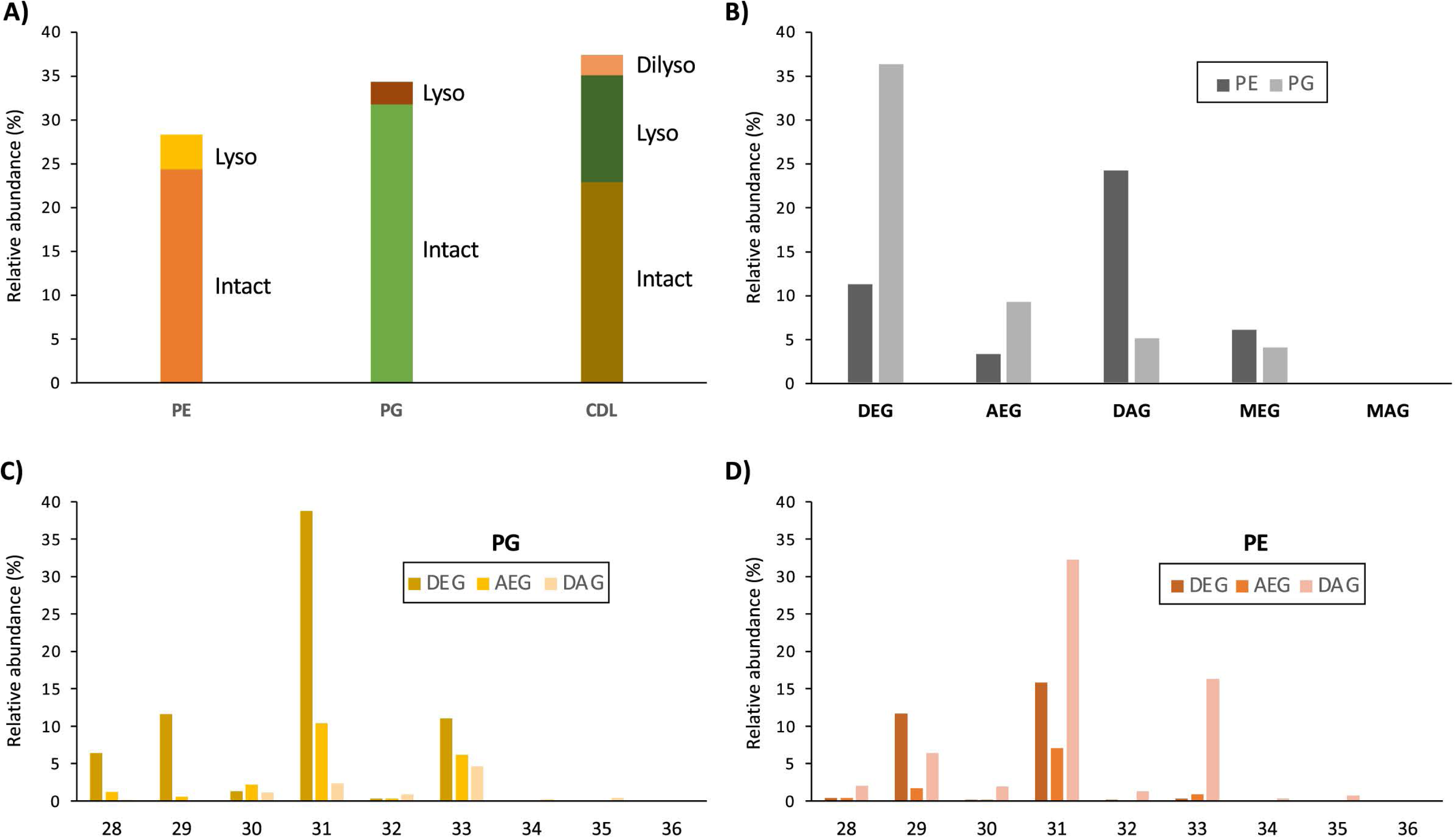
**A)** Relative abundances (% of total membrane phospholipids) of intact and lyso membrane phospholipids (PE = phosphoethanolamine, PG = phosphoglycerol, CDL = cardiolipin) of *D. alkenivorans* grown on C_16_ *n*-alk-1-ene. Relative abundance (%) of **B)** dialkylglycerides (DEG), acyl/alkylglycerides (AEG), diacylglycerides (DAG), mono- alkyglycerides (MEG) and monoacylglycerides (MAG) with a PE or PG polar head group (normalized to total PG+PE); **C)** DEG-, AEG- and DAG-PGs and **D)** DEG-, AEG- and DAG- PEs with increasing number of carbon atoms (sum of the two alkyl/acyl chains).

The distribution of the PEs and PGs with various cores (DEG, AEG, DAG, MEG, and MAG) was determined by peak integration of specific mass chromatograms (see experimental) (Figure 3B). Within PGs, core lipids with at least one ether-bound alkyl chain dominated (ca. 91%), with DEG core lipids contributing 66.2%, AEG core lipids 17.0%, and MEG core lipids 7.4%. Only 9.3% of the PGs contained a DAG core lipid and MAG-PGs were not detected. Ether lipids were less represented (ca. 46%) in PEs, which were dominated by DAGs (54.0%), followed by DEGs (25.0%), MEGs (13.5%) and AEGs (7.5%), and only traces of MAGs (0.2%). Similarities can be observed between the distribution of the total number of alkyl/acyl carbon atoms within the PE and PG phospholipids (Figures 3C-D). For both PGs and PEs this number is predominantly 31, followed by 33 and 29. The two most dominant phospholipids were C_31_ DEG-PG (representing 39% of total PG and 22% of summed PG and PE) and C_31_ DAG-PE (representing 32% of total PE and 14% of summed PG and PE). Fragmentations observed in the MS^2^ spectra of the various PGs and PEs revealed that their core lipids were composed of C_14_ to C_17_ alkyl and acyl chains, in good agreement with the range of total number of alkyl/acyl carbon atoms (28-36).

These findings are in accordance with the previously reported core lipid composition observed after cell hydrolysis of *D. alkenivorans* grown on hexadec-1-ene (Grossi et al., 2015). This study indicated that the major membrane lipids were composed of *n*-C_14,_ *n*-C_16_ and 10-Me-C_16_ fatty acids, *sn*-1-O-10-Me-C_16_ and *sn*-1-O-*n*-C_16_ (and to a lesser extent *sn*-1-O-C_14_, *sn*-2-O-C_14_ and *sn*-1-O-10-Me-C_14_ homologues) glycerol monoethers, and a *sn*-1-O-10-Me-C_16_-*sn*-2-O-C_14_ glycerol diether (accompanied by smaller amounts of *sn*-1-O-C_16_-*sn*-2-O-C_14_, *sn*-1-O-C_14_-*sn*- 2-O-C_14_, *sn*-1-O-10-Me-C_14_-*sn*-2-O-C_16_, *sn*-1-O-10-Me-C_16_-*sn*-2-O-C_16_ homologues). Our UHPLC-ESI/HRMS^n^ method does not allow to identify the degree and position of branching of the alkyl/acyl chains. However, based on the core lipid composition of this strain grown under similar conditions (Grossi et al., 2015), it is likely that i) the alkyl/acyl moieties with 14 and 16 carbons atoms identified in the PE and PG phospholipids are *n*-C_14_ and *n*-C_16_ carbon chains, respectively, ii) those with 15 and 17 carbon atoms identified in the PE and PG phospholipids are 10-Me-C_14_ and 10-Me-C_16_, respectively and, iii) these branched alkyl/acyl moieties are only located at the *sn*-1 position of the glycerol moiety of the phospholipids and not at the *sn*-2 position.

### Identification and Distribution of Alkyl Ether Cardiolipins

In addition to the dialkyl, monoalkyl/monoacyl, and diacyl PE- and PG-phospholipids, we observed series of high molecular weight (MW) phospholipids. Figure 4 shows a typical fragmentation (MS^2^) spectrum of one of these high MW phospholipids eluting at 19.75 min in the BPC (Figure 2), which exhibits a parent ion of *m/z* 1298.063 (exact formula C_73_H_151_O_13_P_2_, Δ ppm −1.6). After an initial loss of H_2_O to produce the fragment ion at *m/z* 1280.050, two separate losses of 526.533 Da (C_34_H_70_O_3_, Δ ppm 1.6) and 554.566 Da (C_36_H_74_O_3_, Δ ppm 3.4) were observed to produce fragments at *m/z* 771.529 (C_39_H_81_O_10_P_2_, Δ ppm −1.6) and *m/z* 743.497 (C_37_H_77_O_10_P_2_, Δ ppm 1.6), respectively. A further loss of 79.967 Da (HPO_3_, Δ ppm 2.6) from each these two fragments produces fragment ions at *m/z* 691.563 (C_39_H_80_O_7_P) and 663.531 (C_37_H_76_O_7_P), respectively. In the lower mass range of the spectrum, a series of fragments at *m/z* 255.268 (C_17_H_35_O, Δ ppm −3.5), *m/z* 283.299 (C_19_H_39_O, Δ ppm −3.1), and *m/z* 297.315 (C_20_H_41_O, Δ ppm −2.8) are observed that appear to be related to the alkyl chains of the phospholipid. Two P-containing fragments at *m/z* 155.010 (C_3_H_8_O_5_P, Δ ppm −3.1) and 234.977 (C_3_H_9_O_8_P_2_, Δ ppm −2.1) likely originate from the head group. Based on this fragmentation pattern, we identify this lipid as a CDL with two different DEG core lipids. The initial loss of H_2_O likely originates from the free hydroxyl of the central, third glycerol moiety of the CDL. The two subsequent losses of 526 and 554 Da represent the loss of a DEG core lipid with 31 and 33 carbon atoms (sum of the two alkyl chains without the glycerol group), respectively. A further loss of a phosphatidic acid from these two fragment ions results in the fragment ions at *m/z* 691 and 665, representing the remaining DEG with a PG moiety. The two phosphor- containing fragments at *m/z* 235 and 155 represent (part of) the central characteristic phosphatidyl-glycero-phosphatidyl (PGP) moiety of CDLs. The fragments at *m/z* 255, 283 and 297 are interpreted to correspond to the different glycerol-bound alkyl moieties contained in the two DEG core lipids. Based on the core lipid composition described above and in a previous study (Grossi et al., 2015), we surmise that the DEG with 31 non-glycerol carbon atoms is composed of a glycerol with ether-bound *n*-C_14_ and 10-Me-C_16_ alkyl moieties at the *sn*-2 and *sn*-1 positions, respectively. The other DEG moiety with 33 non-glycerol carbon atoms consists of a glycerol with ether bound n-C_16_ (*sn*-2) and 10-Me-C_16_ (*sn*-1) alkyl moieties. Together they form a tetraether CDL with 64 alkyl carbon atoms. To the best of our knowledge, this is the first description of a bacterial tetraether CDL with four non-isoprenoidal aliphatic alkyl moieties. Based on this structural elucidation, we further identified a range of tetraether CDLs with the number of alkyl carbon atoms ranging from 56-66, dominated by 62, and, to a lesser extent, 64, 60, and 59 (Figure 5A).

**Figure 4.**
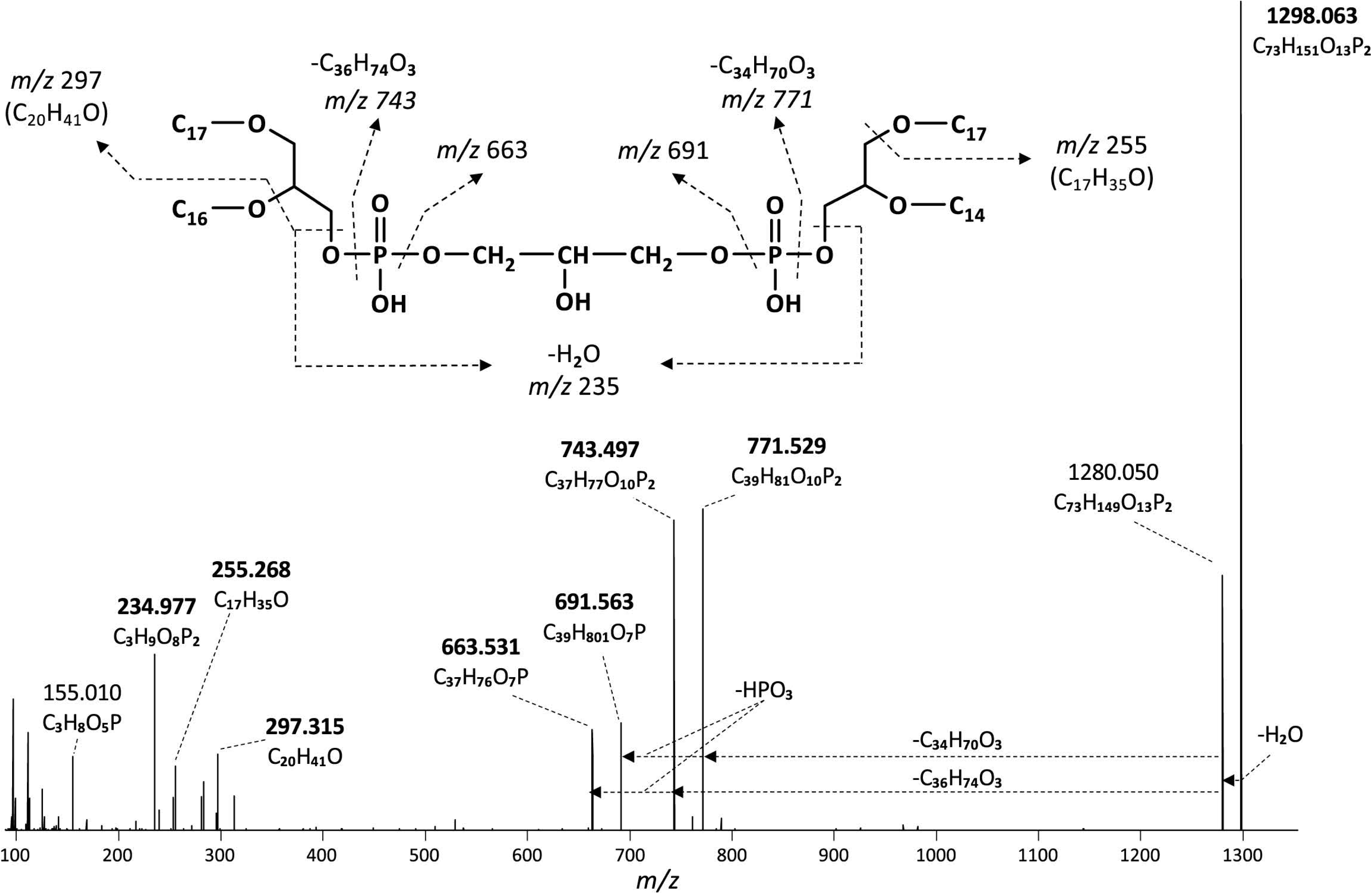
MS^2^ mass spectrum (UHPLC-ESI/HRMS^n^) of m/z 1298.1 ([M+H]^+^) of a tetraether cardiolipin with 64 alkyl carbon atoms (C_33_DEG/C_31_DEG; Figure 6A) synthesized by *D. alkenivorans* grown on C_16_ *n*-alk-1-ene. Based on the GC-MS analysis of hydrolysed core lipids, C_17_ alkyl chains are 10-MeC_16_ carbon chains located at *sn*-1 position, whereas the C_14_ and C_16_ alkyl chains are unbranched *n*-C_14_ and *n*-C_16_.

**Figure 5.**
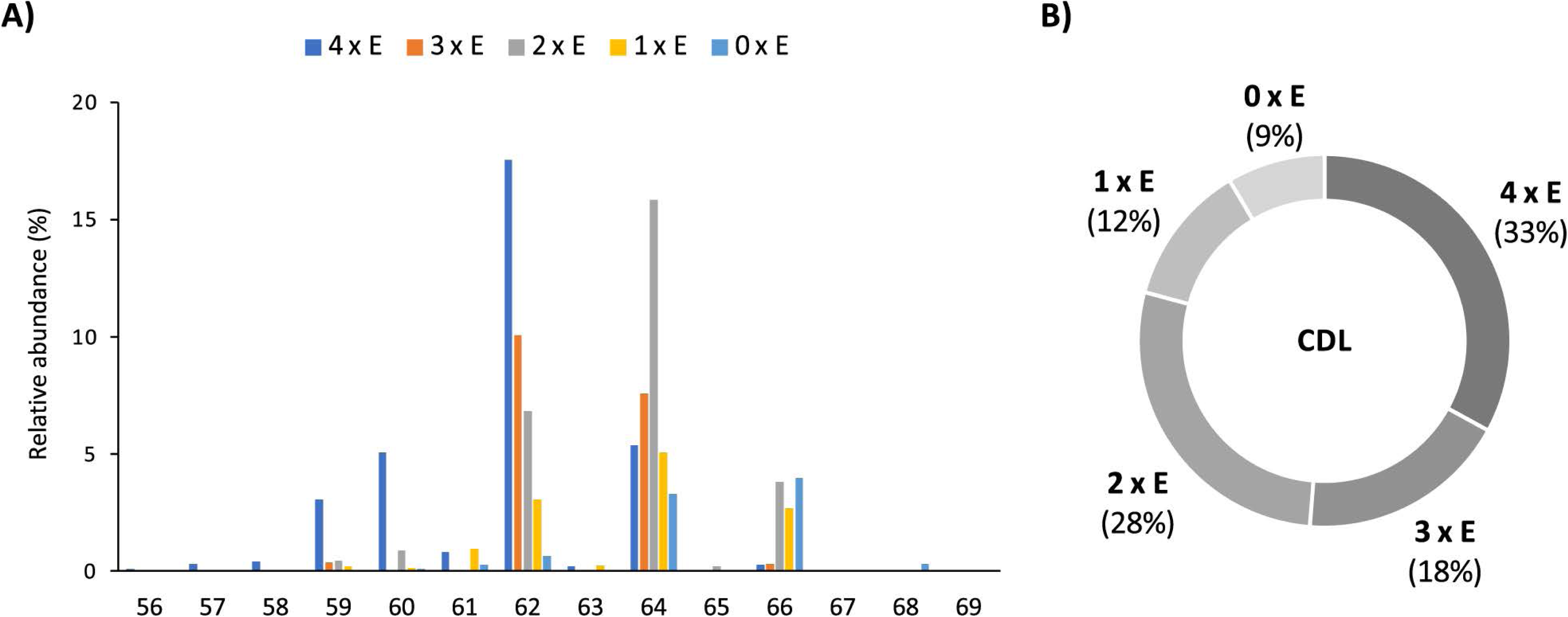
Relative abundances (%) of **A)** tetraethers (4 x E) to tetraesters (0 x E) CDLs with increasing number of carbon atoms (sum of the four alkyl/acyl chains), and **B)** tetraethers (4 x E) to tetraesters (0 x E) among total intact CDLs in *D. alkenivorans* grown on C_16_ *n*-alk-1-ene.

Based on the distribution of core lipids in the PE and PG phospholipids (see previous section), we also searched for additional CDLs containing combinations of DEG, AEG, and DAG cores. A full range of CDLs with 56 to 68 alkyl/acyl carbon atoms in their core lipids and with a number of ether-bound alkyl chains ranging from 0 to 4 (i.e., from tetraesters to tetraethers) was observed in varying relative abundances (Figure 5). Tetraether (33%) and diether/diester (28%) CDLs were most abundant followed by triether/monoester (18%) and monoether/triester (12%), while tetraesters represented only 9% of the total CDLs (Figure 5B). This overall CDL composition of *D. alkenivorans* appears in good agreement with the DEG/AEG/DAG composition observed in PEs and PGs (Figure 3B) and with the DEG/MEG composition of hydrolysed cells of *D. alkenivorans* (Grossi et al., 2015). The high compositional complexity in the distribution of the CDLs is exemplified by the extracted ion chromatograms of the CDLs with 64 alkyl/acyl carbon atoms in their two cores and a decreasing number of ether bonds (Figure 6). An increase in the number of ether alkyl chains in the CDL results in a reduced retention time. The identification of these mixed ether/ester CDLs is supported by their elemental composition, showing one additional oxygen atom for each following member of the series (Figure 6), and their mass spectral fragmentation patterns (Figure 7). Unlike the MS^2^ spectrum of the tetraether CDL (Figure 4), the spectra of CDLs containing one or more acyl groups [i.e., triether/monoester (Figure 7A), diether/diester (Figure 7B), monoether/triester (Figure 7C) and tetraester (Figure 7D)] are characterized by a limited number of fragment ions representing the AEG or DAG core(s). Fragments related to DEG core lipids were low in abundance or not observed. The total number of alkyl/acyl carbons atoms of these moieties were deduced from a neutral loss from the parent ion and from the elemental composition of the compound.

**Figure 6.**
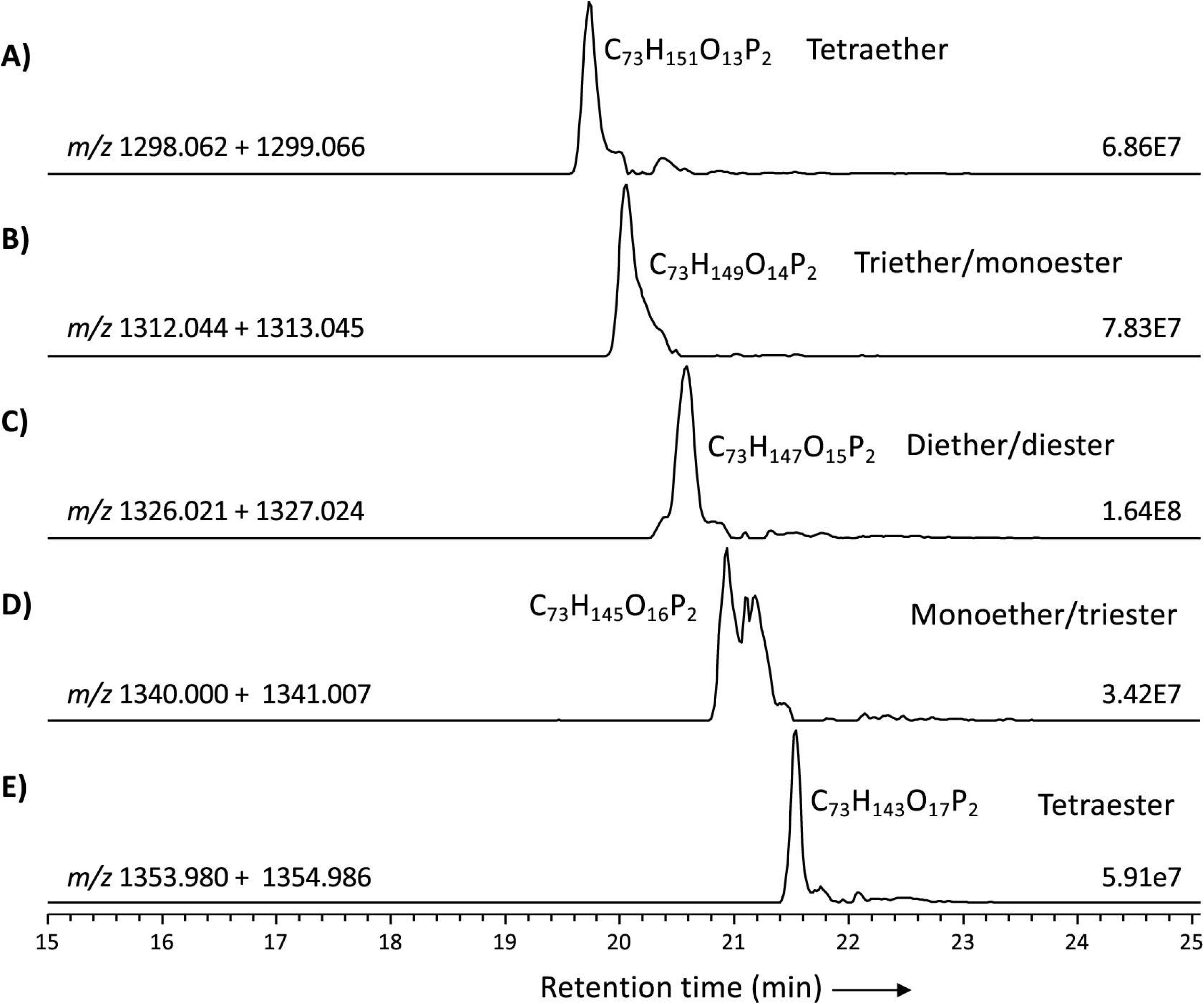
Partial mass chromatograms of CDLs with 64 alkyl/acyl carbon atoms in *D. alkenivorans* grown on C_16_ *n*-alk-1-ene, with decreasing number of ether-bound alkyl chains. Each trace is labeled with the exact mass used for detection, and the intensity of the highest peak is in arbitrary units (AU). MS^2^ spectra are shown in figures 4 and 6.

**Figure 7.**
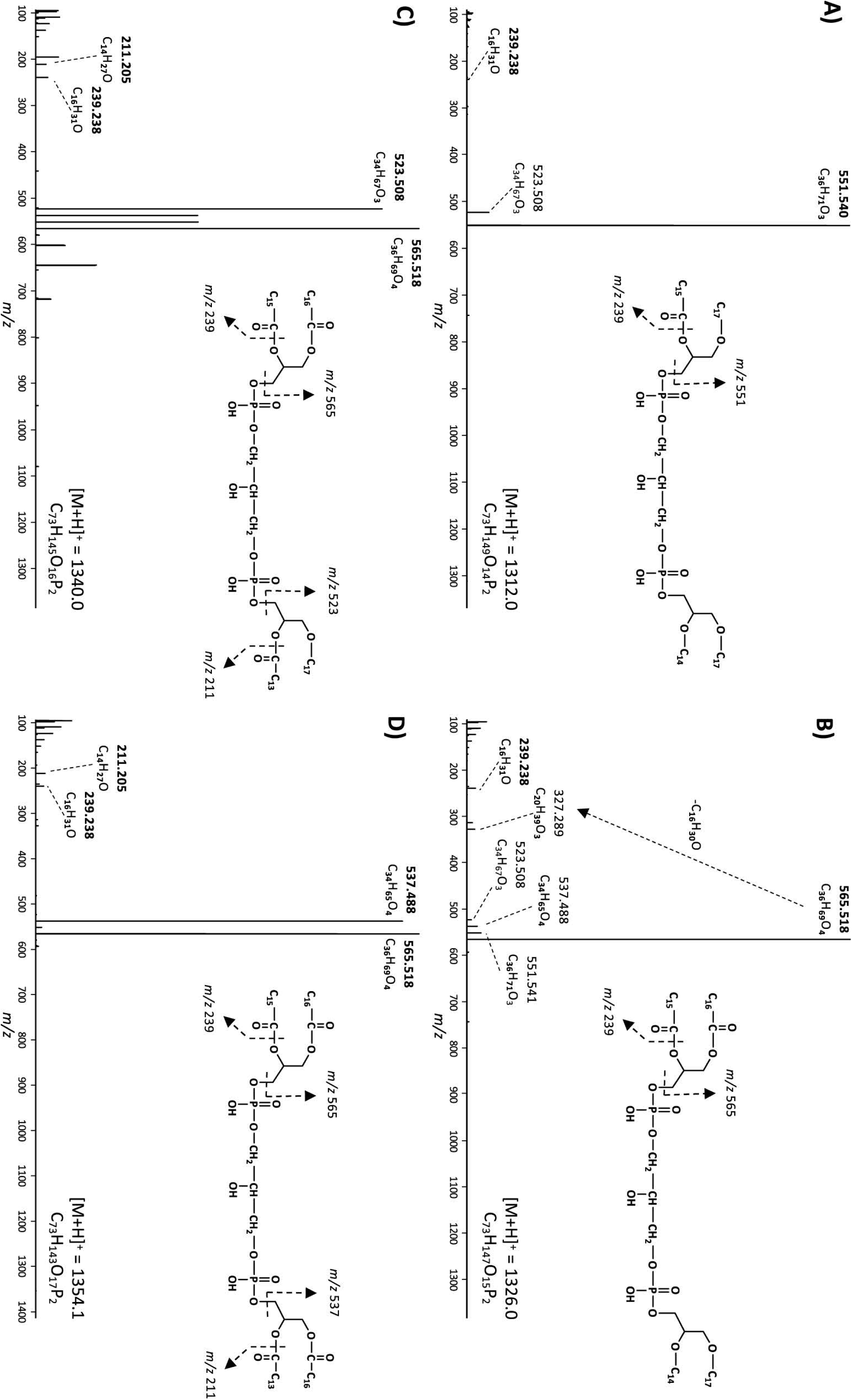
MS^2^ mass spectra (UHPLC-ESI/HRMS^n^) of triether/monoester to tetraester CDLs with 64 alkyl/acyl carbon atoms in *D. alkenivorans* grown on C_16_ n-alk-1-ene. **A)** triether/monoester (m/z 1312.0, [M+H]^+^, Figure 6B) consisting of a C_33_ AEG/C_31_ DEG, co- trapped with a C_33_ DEG/C_31_ AEG; **B)** diether/diester (m/z 1326.0, [M+H]^+^, Figure 6C) consisting of a C_31_ DEG/C_33_ DAG, co-trapped with a C_33_ DEG/C_31_ DAG; **C)** monoether/triester (m/z 1340.0, [M+H]^+^, Figure 6D) consisting of a C_31_AEG/C_33_DAG, co-trapped with a C_31_ DAG/C_33_ AEG; **D)** tetraester (m/z 1354.1, [M+H]^+^, Figure 6E) consisting of a C_33_ DAG/C_31_ DAG. Based on the GC-MS analysis of hydrolysed core lipids, C_17_ and C_15_ alkyl/acyl chains are 10-MeC_16_ and 10-MeC_14_ carbon chains located at *sn*-1 position, whereas the C_14_ and C_16_ alkyl/acyl chains are unbranched *n*-C_14_ and *n*-C_16_ carbon chains.

In most classes of CDLs, components with 62 or 64 alkyl/acyl carbon atoms were most abundant (Figure 5A), except for the tetraester CDLs, where the member with 66 acyl carbon atoms was most dominant. Overall, the tetraether with 62 alkyl/acyl carbon atoms (18%) and the diether/diester with 64 alkyl/acyl carbon atoms (16%) were the two most dominant compounds among total CDLs, followed by the triether/monoester with 62 or 64 alkyl/acyl carbon atoms.

### Identification and Distribution of Lyso-cardiolipins

In addition to the above described tetraether CDLs, we detected several series of ether alkyl CDLs lacking one or two acyl chains, i.e. so-called monolyso- and dilyso- ether alkyl CDLs, respectively (Figure 1). Trilyso-CDLs were not detected. Figure 8 shows diagnostic MS^2^ spectra of a triether monolyso-CDL and of a diether dilyso-CDL with a total number of alkyl carbon atoms of 45 and 31, respectively. As for the tetraether CDLs, their mass spectra show specific fragmentations corresponding to the loss of one of the two alkylglycerol cores. The intensity of these specific fragment ions and of the parent ion decreases, however, as the number of ether-linked alkyl chains decreases (Figures 4 and 8).

**Figure 8.**
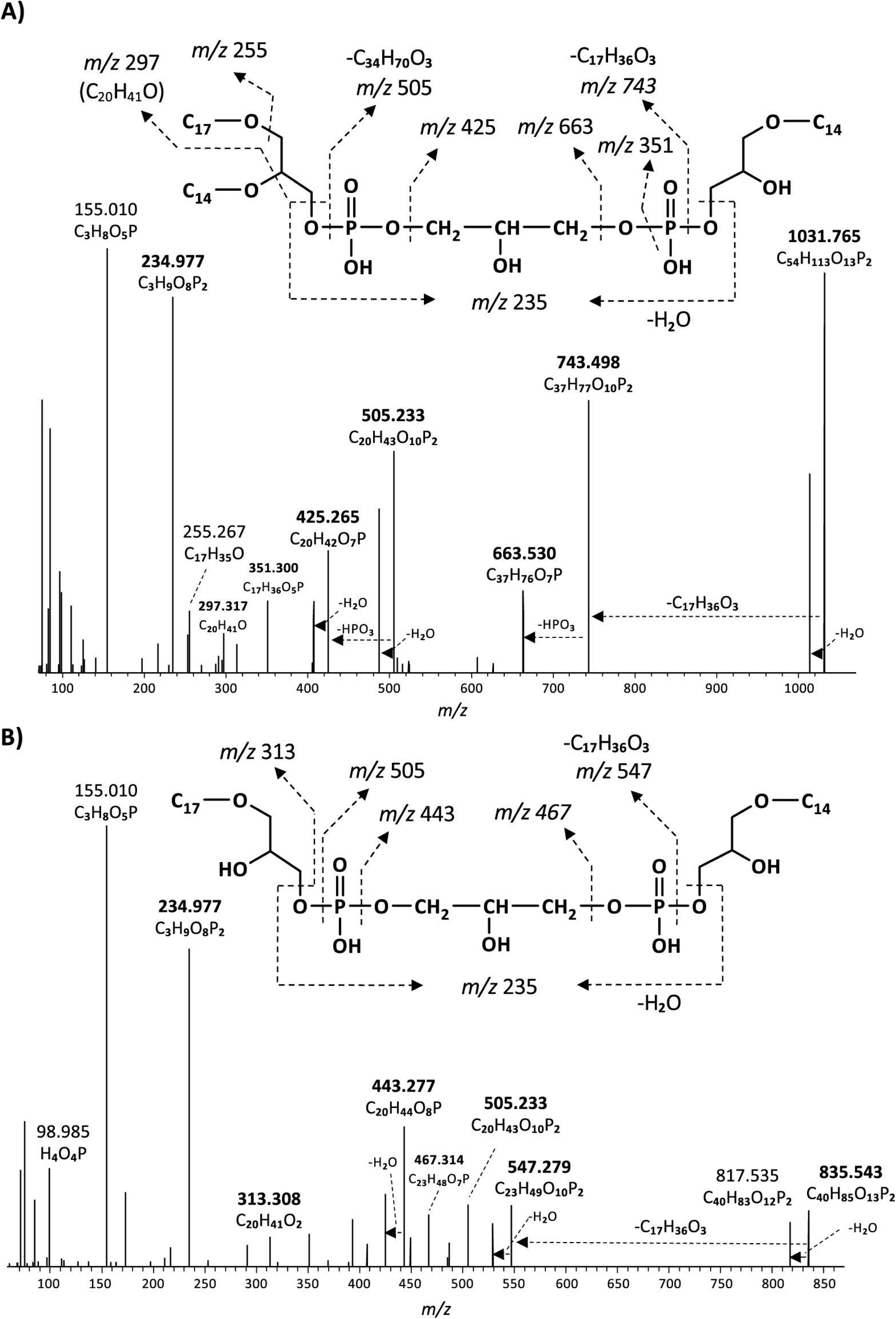
MS^2^ mass spectra (UHPLC-ESI/HRMS^n^) of **A)** m/z 1031.8 ([M+H]^+^) of a monolyso- cardiolipin with 45 alkyl carbon atoms and 3 ether bound alkyl chains composed of a C_14_MEG and a C_31_DEG containing C_14_ and C_17_ alkyl chains (Figure 9A), and **B)** m/z 835.5 ([M+H]^+^) of a dilyso-cardiolipin with 31 alkyl carbon atoms and 2 ether bound alkyl chains composed of a C_14_MEG and a C_17_MEG (Figure 9B). Based on the GC-MS analysis of hydrolysed core lipids, the C_17_ alkyl chains are 10-MeC_16_ carbon chains located at *sn*-1 position, and the C_14_ alkyl chain is an unbranched *n*-C_14_.

Like intact CDLs, lyso-CDLs also occurred as series of mixed ether/ester compounds, ranging from triethers to triesters and from diethers to diesters for monolyso- and dilyso-CDLs, respectively (Figures 9 and S1). As for intact CDLs, an increasing number of ether alkyl chains in the lyso-CDLs results in a decreased retention time. Dilyso-CDLs were observed in two structural variations: either two monolyso core lipids (i.e., MEG/MEG or MEG/MAG) connected by a PGP moiety, or an intact core lipid (with two alkyl/acyl chains) linked to a PGPG head group (Figure 9). Mono-lyso CDLs ranged from C_42_ to C_52_ (sum of the three alkyl/acyl chains) and were dominated by a C_47_ monoether/diester and C_45_ and C_48_ triethers (Figure 10A), whereas di-lyso CDLs ranged from C_28_ to C_35_ (sum of the two alkyl/acyl chains) and were dominated by a C_31_ diether (Figure 10B).

**Figure 9.**
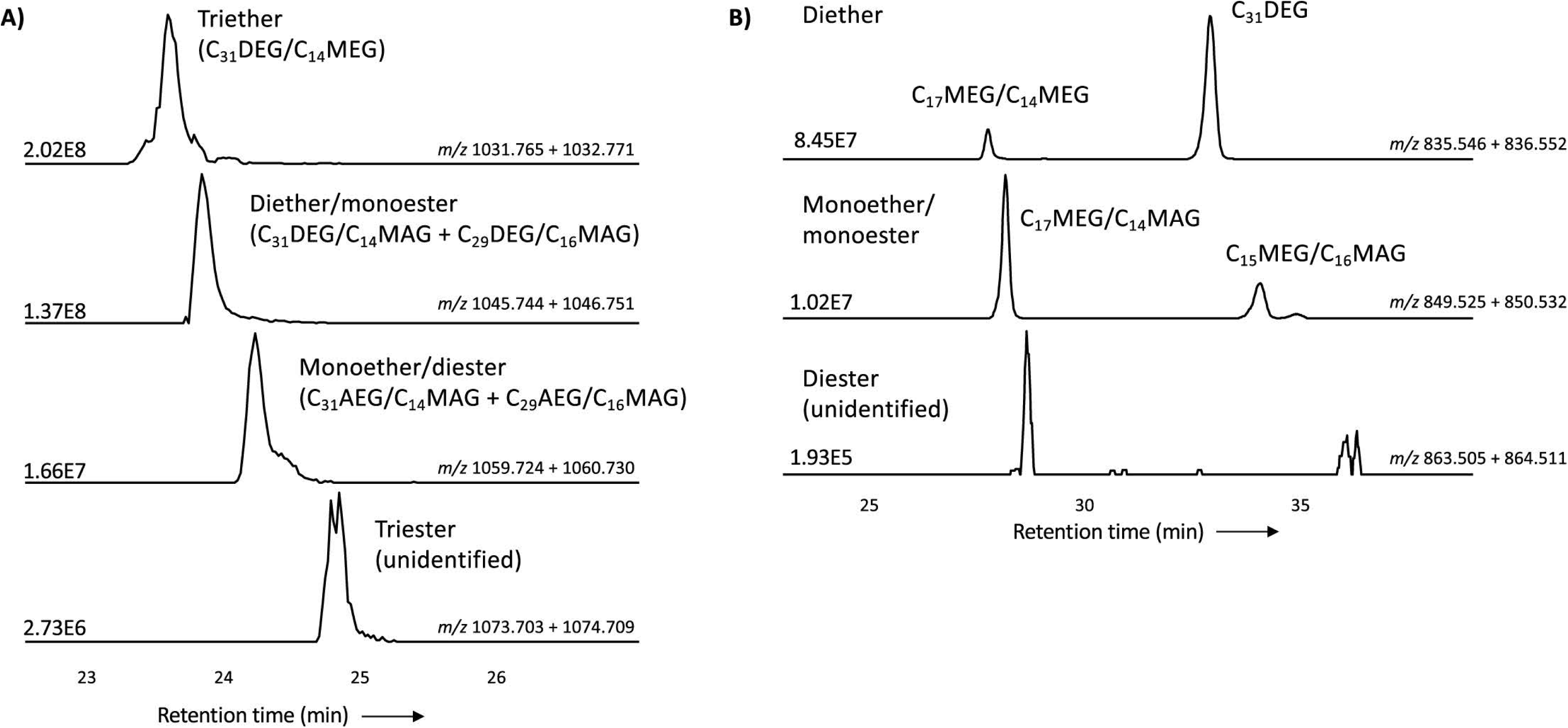
Partial mass chromatograms of **A)** monolyso-CDLs with 45 alkyl/acyl carbon atoms and 3 - 0 ether bound alkyl chains and **B)** dilyso-CDLs with 31 alkyl/acyl carbon atoms and 2 - 0 ether bound alkyl chains in *D. alkenivorans* grown on C_16_ *n*-alk-1-ene. Each trace is labeled with the exact mass used for detection, and the intensity of the highest peak is in arbitrary units (AU). The strong fragmentation of triester monolyso-CDLs and diester dilyso-CDLs did not allow determination of their exact structures based on MS^2^ mass spectra. DEG = dialkylglyceride, AEG = acyl/alkylglyceride, MEG = mono-alkyglyceride, MAG = monoacylglyceride. Unidentified = unavailability of MS^2^ mass spectra.

**Figure 10.**
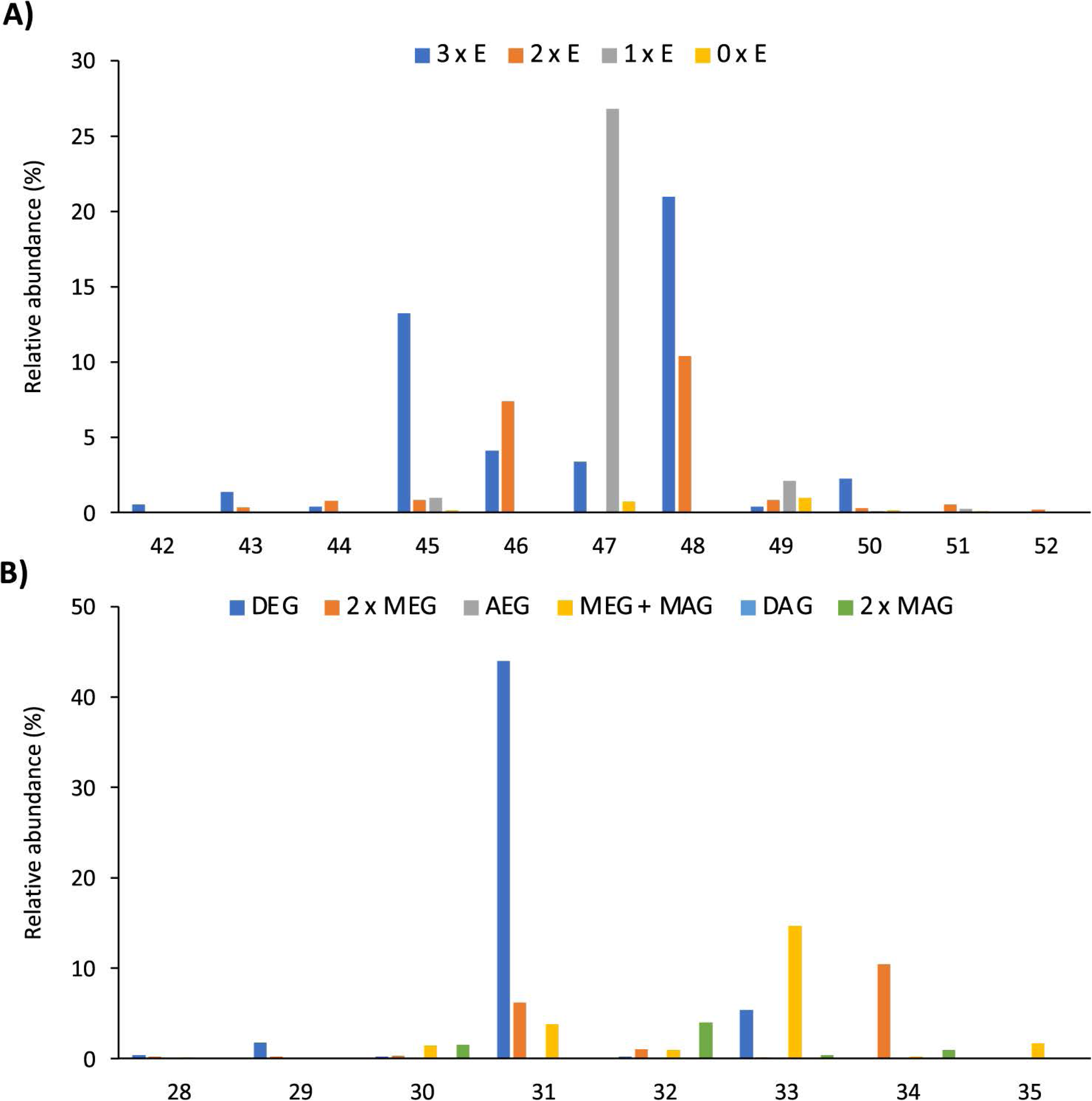
Distributions (%) of number of carbon atoms (sum of the three or two alkyl/acyl chains) of **A)** all monolyso-CDLs (from triethers (3 x E) to triesters (0 x E)) and **B)** all dilyso- CDLs in *D. alkenivorans* grown on *n*-hexadec-1-ene. DEG = dialkylglyceride, AEG = acyl/alkylglyceride, DAG = diacylglyceride, MEG = mono-alkyglyceride, MAG = monoacylglyceride.

Monolyso-CDLs (ca. 12% in total. Fig. 3a) were much more abundant than dilyso-CDLs (2%) but were present in lower abundance compared to intact membrane lipids represented by CDLs (23%), PGs (32%) and PEs (24%). However, lyso-components form a much larger fraction of the total CDLs than in case of the PEs and PGs (i.e. 38% vs. 14 and 9%, respectively).

### Potential Enzymes Involved in the Biosynthesis of Tetraester/Tetraether CDLs

Potential enzymes involved in the synthesis of CDLs in *D. alkenivorans* were identified by homology search (pBLAST) using the protein sequences of the three CDL synthases ClsA, ClsB, and ClsC of *E. coli* (Tan et al., 2012) as queries. This search resulted in three hits, i.e. with the proteins SHL34086, SHI83627 and SHL08500 (Table S1-2), which were also found to be annotated as CDL synthases in the Uniprot database (Uniprot, 2023) and also classified as members of the PLD superfamily of proteins. Indeed, Cls1 (SHL34086), Cls2 (SHI83627), and Cls3 (SHL08500) contain conserved amino acids necessary for the predicted enzymatic activity when compared to characterized Cls enzymes belonging to the PLD family (Figures S2-3). Hence, our search identified three different potential CDL synthases in *D. alkenivorans*: Cls1, Cls2, and Cls3 (Table S1). Sequence analysis of the predicted Cls suggests Cls1 possesses two transmembrane domains, as predicted for the ClsA of *E. coli* (Tan et al., 2012). Cls1 is also phylogenetically most closely related to the *E. coli* ClsA sequence and the archaeal Cls of *Methanospirillum hungatei* (Exterkate et al., 2021) as shown in the unrooted phylogenetic tree (Figure 11) and sequence comparisons (Table S2). On the other hand, the closely related sequences of Cls2 and Cls3 are arranged together in a separate cluster of Cls (Figure 11) and are not closely related to the ClsB and ClsC proteins of *E. coli* (Tan et al., 2012). Remarkably, Cls3 has no transmembrane domains, while Cls2 only possesses one domain in contrast to Cls1, which contains two domains, like ClsA of of *E. coli* (Tan et al., 2012). pBLAST searches of the genome of *D. aliphaticivorans*, a related species to *D. alkenivorans*, revealed a similar set of three Cls proteins (Table S2). Lastly, the sequence of an eukaryotic-like CDL synthase identified in *Streptomyces coelicolor* A3 (Sandoval-Calderon et al., 2009; Sco1389), which belongs to the CDP-alcohol phosphatidyltransferase superfamily, was also used in a pBLAST search. This resulted in a poor fit with a putative CDP-diacylglycerol-glycerol-3-phosphate, 3- phosphatidyl–transferase protein (SHK52233; Table S2), suggesting that this pathway for CDP synthesis is likely not occurring in *D. alkenivorans*.

**Figure 11.**
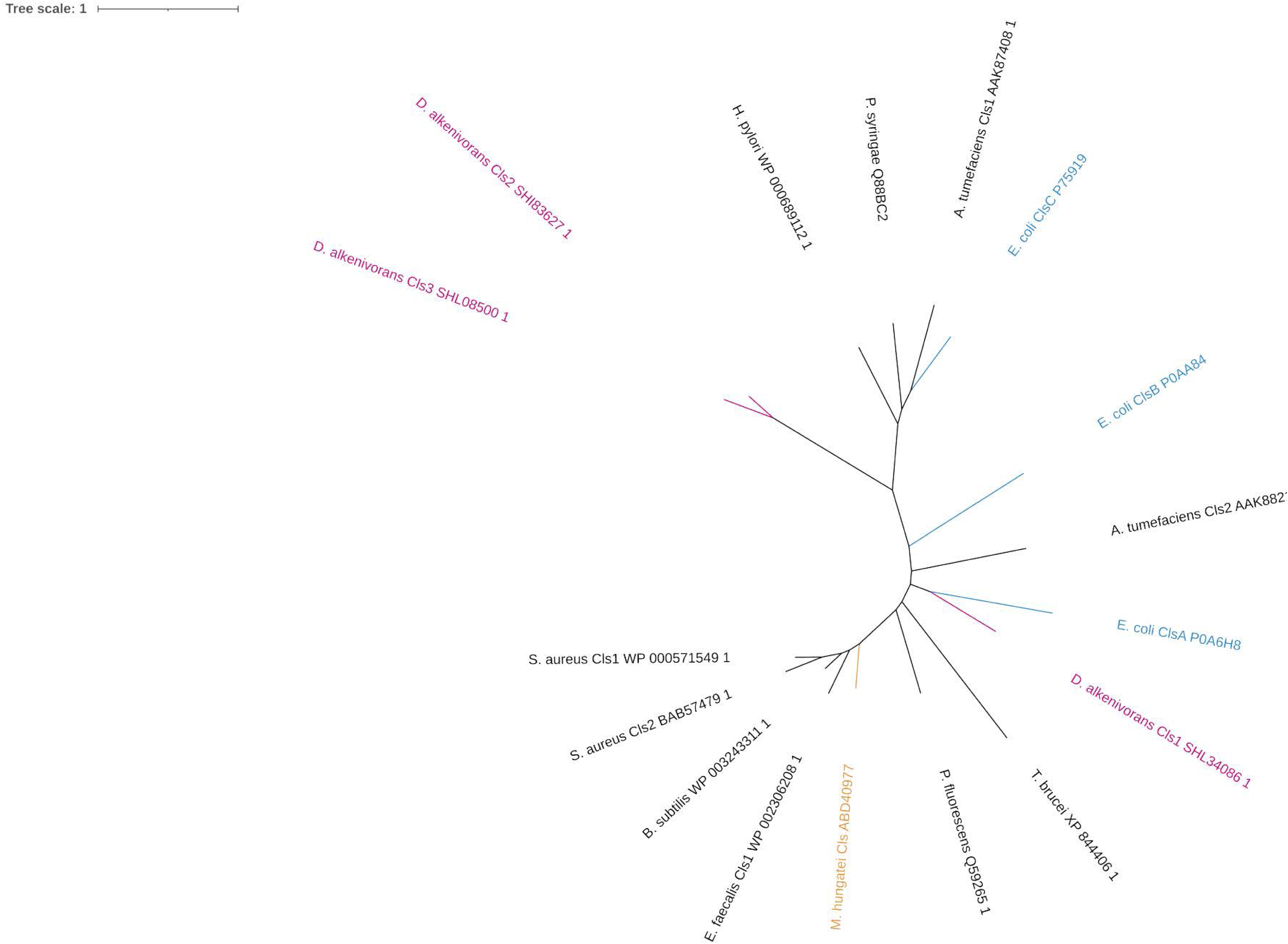
Unrooted tree of putative Cls of *D. alkenivorans* with characterized Cls in bacterial species, in one archaeal species, and in one eukaryotic sequence proposed to possess the same enzymatic activity. The Cls of *D. alkenivorans* are highlighted in pink. Cls 1 is located close to ClsA of *E. coli*, while Cls2 and Cls3 are found together next to other bacterial Cls in a separate cluster.

In an attempt to identify the enzyme responsible for the formation of the alkyl ether bond at the *sn*-2 position of phospholipids (including CDLs) of D. *alkenivorans*, a pBLAST search of its genome using the Ger enzyme (SHJ90043), previously shown to be responsible for the formation of *sn*-1 alkyl ether bonds in glycerol membrane lipids in bacteria (Sahonero-Canavesi et al., 2022), was performed and resulted in the detection of a closely related protein (SHK01260; Tables S1-2). To test the functionality of this enzyme, we performed both aerobic and anaerobic heterologous gene-expression experiments in *E. coli* using the gene itself and a combination with the Ger protein of *D. alkenivorans* (Table 1) using the same methodology as described previously (Sahonero-Canavesi et al., 2022). In the experiments without Ger we did not detect production of glycerol ethers (Table 1). Surprisingly, however, in contrast to earlier findings (Sahonero-Canavesi et al., 2022), we detected in addition to the sn-1 glycerol monoethers, small amounts of three glycerol diethers (C_28:0_, C_30:0_, and C_34:1_; Table 2) in the base-hydrolysed phospholipid extract in the experiment when only the Ger enzyme was expressed (Table 1). This is probably because we analyzed the membrane lipids produced by UHPLC-HRMS^n^, which has a lower detection limit. In the gene expression experiment where Ger was combined with the Ger homologue, glycerol diethers were also detected (Table 2). However, their relative abundance did not increase. Hence, our heterologous gene expression analysis suggests that only the Ger enzyme seems required for the transformation of a glycerol diester into a glycerol diether under the conditions tested here. It is, therefore, responsible for making *D. alkenivorans* capable of converting glycerol ester bonds into ether bonds at both the *sn*-1 and *sn*-2 positions, explaining the presence of DEG, AEG, and DAG cores in PE and PG phospholipids and in the novel CDLs.

**Table 1.** Formation of mono- and dialkyl glycerol ethers after heterologous gene-expression of enzymes of *D. alkenivorans* potentially involved in ether bond formation in *E. coli* BL21 DE3.

| Experiment | Monoethers<sup>a</sup> |  |  |  | Diethers<sup>b</sup> |  |  |
| --- | --- | --- | --- | --- | --- | --- | --- |
|  | C<sub>14:0</sub> 1-*O*-<br>MEG | C<sub>16:0</sub> 1-*O*-<br>MEG | CyC<sub>17</sub> 1-*O*-<br>MEG | C<sub>18:1</sub> 1-*O*-<br>MEG | C<sub>28:0</sub> DAG | C<sub>30:0</sub> DAG | C<sub>34:1</sub> DAG |
| pET29b (negative control) | - | - | - | - | - | - | - |
| Ger (9SHJ90043) | + | + | + | + | + | + | + |
| Ger-like (SHK01260) | - | - | - | - | - | - | - |
| Ger + Ger-like | + | + | - | - | + | + | - |

Key: + = detected; - = not detected; <sup>a</sup> monoethers were detected based on GC-MS analysis after base hydrolysis of the Bligh-Dyer extract; <sup>b</sup> diethers were detected by UHPLC-HRMS<sup>n</sup> analysis after base hydrolysis of the Bligh-Dyer extract.

**Table 2.**
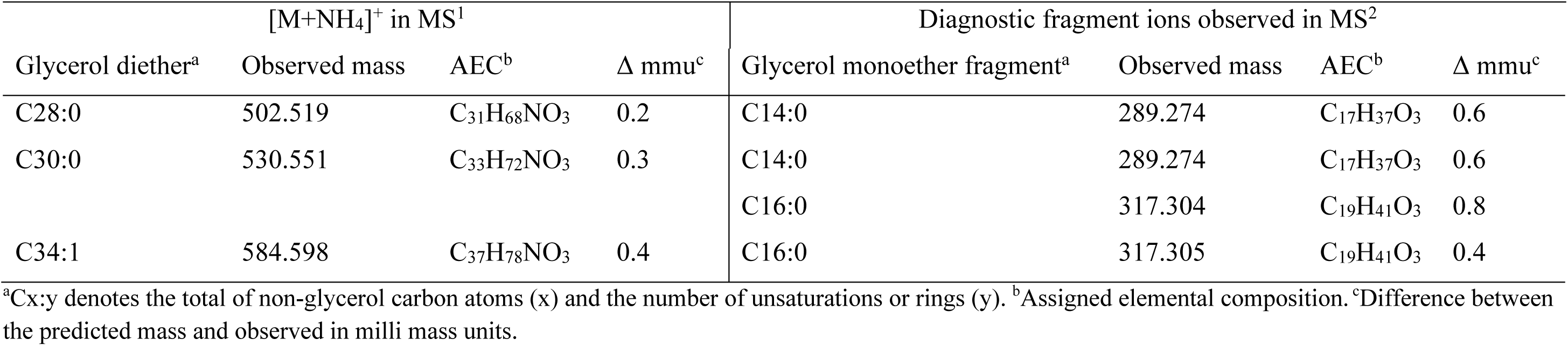
Mass spectral characteristics of diethers detected upon heterologous gene-expression of the Ger enzyme of *D. alkenivorans* in *E. coli* BL21 DE3.

| [M+NH<sub>4</sub>]<sup>+</sup> in MS<sup>1</sup> |  |  |  | Diagnostic fragment ions observed in MS<sup>2</sup> |  |  |  |
| --- | --- | --- | --- | --- | --- | --- | --- |
| Glycerol diether<sup>a</sup> | Observed mass | AEC<sup>b</sup> | Δ mmu<sup>c</sup> | Glycerol monoether fragment<sup>a</sup> | Observed mass | AEC<sup>b</sup> | Δ mmu<sup>c</sup> |
| C28:0 | 502.519 | C<sub>31</sub>H<sub>68</sub>NO<sub>3</sub> | 0.2 | C14:0 | 289.274 | C<sub>17</sub>H<sub>37</sub>O<sub>3</sub> | 0.6 |
| C30:0 | 530.551 | C<sub>33</sub>H<sub>72</sub>NO<sub>3</sub> | 0.3 | C14:0 | 289.274 | C<sub>17</sub>H<sub>37</sub>O<sub>3</sub> | 0.6 |
|  |  |  |  | C16:0 | 317.304 | C<sub>19</sub>H<sub>41</sub>O<sub>3</sub> | 0.8 |
| C34:1 | 584.598 | C<sub>37</sub>H<sub>78</sub>NO<sub>3</sub> | 0.4 | C16:0 | 317.305 | C<sub>19</sub>H<sub>41</sub>O<sub>3</sub> | 0.4 |

<sup>a</sup>Cx:y denotes the total of non-glycerol carbon atoms (x) and the number of unsaturations or rings (y). <sup>b</sup>Assigned elemental composition. <sup>c</sup>Difference between the predicted mass and observed in milli mass units.

## DISCUSSION

### The unusual phospholipid composition of *D. alkenivorans*

The core lipid composition of the mesophilic sulfate-reducing bacterium *D. alkenivorans* strain PF2803^T^ analysed by GC-MS after acid hydrolysis of cells has previously been shown to contain high proportions of dialkyl- and monoalkyl-glycerol ether core lipids (Grossi et al., 2015; Vinçon-Laugier et al., 2016), a feature previously thought to be the hallmark of (hyper)thermophilic bacteria. The present analysis by UHPLC-HRMS^n^ of the phospholipid composition of *D. alkenivorans* confirms that alkylglycerol ether lipids are the dominant membrane lipids of this species, where they occur as dialkyl- and monoalkyl/monoacyl- glycerols with a PE or PG head group in relatively high abundance (Figures 3A-B). PE and PG phospholipids have been reported as common phospholipids in sulfate-reducing bacteria (e.g., Rütters et al., 2001; Sturt et al., 2004).

Our detailed lipidome analysis of *D. alkenivorans* further demonstrates the presence of strong proportions of CDLs (ca. one third of total phospholipids; Figure 3A) dominated (>90%) by core lipids containing one to four ether bound alkyl chains, and essentially represented by tetraether CDLs and CDLs composed of two mixed alkyl/acyl glycerols (each representing ca. one third of total CDLs). Alkyl ether CDLs thus constitute the third major class of phospholipids of *D. alkenivorans* together with alkyl ether PEs and PGs, which contrasts with the general view that CDLs are minor compounds of bacterial membranes (Schlame, 2008). CDLs with alkyl vinyl ethers (so-called plasmalogens) occur quite commonly in eukaryotes (Goldfine, 2010) and bacteria (e.g., *Clostridium innocuum*; Johnston et al., 1994) but their saturated counterparts are much more seldom. Sturt et al. (2004) suggested their presence in the mesophilic and thermophilic sulfate-reducing bacteria *Desulfosarcina variabilis* and *Thermodesulfobacterium commune* but the technique applied (HPLC/electrospray ionization ion-trap MS^n^) did not allow the structural characterization of CDLs with mixed ether- and ester-linkages. This is now possible by the application of HRMS, as demonstrated here. The present work further revealed the occurrence of non-isoprenoid tetraether CDLs, which is unprecedented. Until now, naturally occurring tetraether CDLs have been reported in methanogens and extremely halophilic archaea (Lattanzio et al., 2002; Sprott et al., 2003; Angelini et al., 2012; Bale et al., 2019) but these contain ether-bound C_20_ and C_25_ isoprenoid alkyl chains.

### Biosynthesis of Alkyl Ether CDLs

In eukaryotes and a few bacteria (Sandoval-Calderon et al., 2018) CDLs are biosynthesized by condensation of PG and cytidine diphosphate-diacylglycerol (CDP-DAG), leading to two phosphatidyl moieties linked by a central glycerol molecule. In contrast, in almost all bacteria, CDL formation proceeds via the condensation of two PG or one PG and one PE molecules (Tan et al., 2012). The enzyme required for the condensation of two PG molecules is also encoded in the genomes of some archaea (Exterkate et al., 2021). Detailed investigation of *E. coli* revealed that its genome encodes three Cls enzymes that catalyze formation of CDLs. ClsA and ClsB catalyze the condensation of two PG molecules, while ClsC mediates the condensation of PG and PE (Tan et al., 2012). For this latter reaction, the presence of another protein, YmdB, substantially increases the production of CDLs (Tan et al., 2012).

In the *D. alkenivorans* genome, no homologue to Sco1389, encoding the “eukaryotic” biosynthetic pathway for CDL production (Sandoval-Calderon et al., 2009), was detected, so formation of CDLs through condensation of PG and CDP-DAG is unlikely. However, we detected three genes encoding for potential Cls enzymes that belong to the PLD family of proteins. All three possessed the two characteristic HKD-motifs present in fully characterized Cls of other bacteria (Figure S2-3). Thus, all three enzymes may be involved in the CDL biosynthetic process in this sulfate-reducing bacterium, like in *E. coli* (Tan et al., 2012). Notably, the sequence and phylogenetic position of Cls1 of *D. alkenivorans* closely resembles that of the characterized ClsA from *E. coli* (Figure 11, Supplementary Table S2) (Quigley and Tropp, 2009). At the same time, the sequence of Cls1 possesses structural and hydrophobic characteristics that are close to the recently characterized archaeal Cls from *Methanospirillum hungatei* (Exterkate et al., 2021) (Table S2). The enzymatic characterization of the *M. hungatei* Cls revealed the promiscuity of this enzyme with respect to the phospholipid species utilized as substrate; it was shown to be non-selective towards the stereochemistry of the glycerol backbone, the nature of the lipid tail or the headgroup, and mode of bonding of the lipid tail (i.e., ester vs. ether bound) (Exterkate et al., 2021). Hence, this enzyme was able to catalyze the biosynthesis of archaeal, bacterial, and mixed archaeal/bacterial CDL species from a wide variety of substrates. The close resemblance of Cls1 of *D. alkenivorans* to ClsA of *E. coli* and Cls of *M. hungatei* indicates that it is likely that Cls1 is able to utilize a wide variety of PG substrates, i.e. with a different core composition (DAG, AEG and DEG), or perhaps even in its lyso forms (MEG and MAG), as biosynthetic precursors for CDL synthesis. The role of Cls2 and Cls3 of *D. alkenivorans*, which are phylogenetically closely related, is less clear-cut. They are less closely related to any of the three Cls enzymes of *E. coli* than Cls1 (Table S2). Phylogenetically, however, they are somewhat related to ClsC of *E. coli* (Figure 11). The *ymdB* gene is also present in the genome of *D. alkenivorans* (Table S1). This supports that Cls2 and/or Cls3 are catalyzing a similar reaction as ClsC in *E. coli*, i.e. condensation of a PE and a PG molecule to form CDL, since the YmdB protein enhances this reaction (Tan et al., 2012). Hence, like in *E. coli, D. alkenivorans* is probably capable of producing CDLs in two ways, by condensation of two PGs and by condensation of a PG and a PE. Interestingly, our analysis revealed a remarkable strong difference between PEs and PGs with respect to the presence of the DAG, AEG, and DEG core lipids; PGs contain relatively more ether bonds in their core than PEs (Figure 3B). The two pathways for the formation of CDLs would thus result in a different composition in terms of the abundance of ether bonds and we can theoretically calculate this, assuming that ester bonds in the core lipids of CDLs are not changed (i.e., reduced to ether bonds) after their biosynthesis. The results of these calculations are shown in Figure 12. The hypothetical condensation of two PE molecules would result in CDLs with a distribution dominated by tetraesters and diesters/diethers (Figure 12a) while condensation of two PG molecules, likely catalyzed by Cls1, would result in CDLs where tetraethers form ca. 50% (Figure 12c). The combination of one PE and PG, as hypothesized to be catalyzed by Cls2 and/or Cls3, would result in CDLs dominated (ca. 50%) by diesters/diethers (Figure 12b). When the measured composition of the CDLs of *D. alkenivorans* (Figure 12e) is considered, no good match with the three calculated CDL distributions (Figure 12a-c) is observed. However, when we assume that both pathways of CDL synthesis, i.e., condensation of two PG molecules and of a PG and a PE molecule, operates similarly at the given growth phase, a CDL distribution is obtained (Figure 12d) that fits quite well with the actual CDL distribution (Figure 12e), i.e. with equal amounts (ca. 30%) of diesters/diethers and tetraethers. This exercise suggests that both biosynthetic pathways for production of CDLs in *D. alkenivorans* are active.

**Figure 12.**
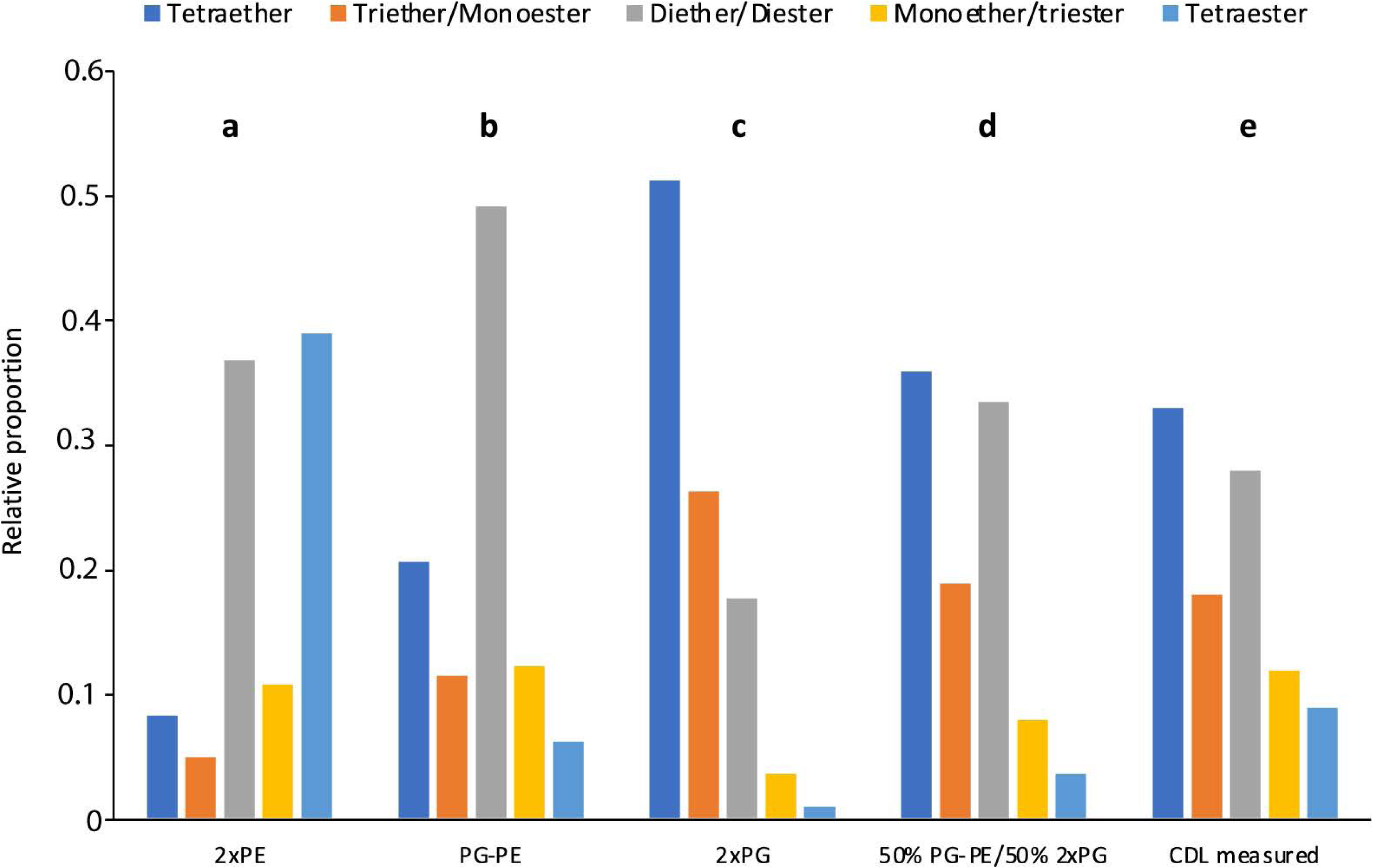
Estimated (**a** – **d**) CDL composition in comparison with the measured CDL composition (**e**) in *D. alkenivorans* grown on *n*-hexadec-1-ene. The estimated CDL composition was calculated from the measured PE and PG composition (Figure 3) assuming that CDLS were formed by condensation of (**a**) two PEs, (**b**) one PE and one PG, (**c**) two PGs, and (**d**) a 50-50 combination of scenario **b** and **c**.

To confirm this hypothesis the role of the three Cls enzymes of *D. alkenivorans* should be tested in its own genomic context. Unfortunately, no genetic tools are currently available to inactivate these genes individually and test how this would affect the abundance and composition of the CDLs. Alternatively, if changes in the cellular content and composition of CDLs could be observed under different growth conditions (e.g., physiological state, pH, growth phase, temperature, nutrient concentration), the expression of individual *cls* genes could be detected and measured by transcriptomic or other gene-expression analysis. This may provide a deeper understanding of these unusual ether lipids and their significance for the physiology of sulfate-reducing bacteria.

The biosynthetic link between CDLs and the two other classes of phospholipids in *D. alkenivorans* (i.e. PEs and PGs; Figure 3A) is further supported by the similar distribution of the alkyl chains present in alkyl ether CDLs, PEs and PGs, which are formed from *n*-hexadec- 1-ene. Since the alkylglycerol ether lipid composition of this heterotrophic sulfate-reducing bacterium has been shown to strongly depend on growth substrate (Vinçon-Laugier et al., 2016), it is expected that variations in carbon source will directly impact the composition of biosynthesized CDLs.

### Insights into alkyl ether lipid biosynthesis

The biosynthesis of *sn*-1 monoalkylglycerols (1-*O*-MEGs) has been confirmed to be catalyzed by Ger, which is a homolog of the plasmalogen biosynthetic enzyme PlsA (Sahonero-Canavesi et al., 2022). Heterologous-gene expression in *E. coli* of the *D. alkenivorans* Ger enzyme confirmed that the main lipid products were 1-*O*-MEGs with ether-bound alkyl chains derived from the fatty acids produced by *E. coli*, in line with previous investigation of the biosynthetic pathways of MEGs and DEGs in *D. alkenivorans* based on labeling experiments (Grossi et al., 2015). Lipids with an ether bond at the *sn*-2 position, i.e., *sn*-2 monoalkylglycerols (2-*O*-MEGs) and *sn*-1,2 dialkylglycerols (DEGs), are also biosynthesized by *D. alkenivorans*, although 2-*O*- MEGs occur in much smaller relative abundances (Grossi et al., 2015; Vinçon-Laugier et al., 2017). Glycerol derivatives with an ether bound alkyl group at the *sn*-2 position were not detected in previous heterologous-gene expression experiments with Ger (Sahonero-Canavesi et al., 2022). This suggested that another enzyme would be responsible for their synthesis.

Individual expression of the suspected gene of *D. alkenivorans* for *sn*-2 ether bond formation, i.e. Ger2, did, however, not yield *sn*-2 ether lipids; in fact no ether lipids were produced at all (Table 1). It remains possible that the Ger2 and or other additional enzymes are involved in ether lipid formation, but that they remained inactive due to the expression conditions in our experimental set-up or, alternatively, a required additional co-factor was absent in the expression host. Genetic manipulation or testing different expression hosts and/or conditions may provide further insight into whether Ger2 is functional as an ether lipid-forming enzyme. Since our previous experiments on testing the enzymatic activity of various Ger proteins often resulted in the accumulation of the protein as inclusion bodies, thus producing inactive versions of the Ger enzymes (Sahonero-Canavesi et al., 2022), we also repeated the experiment with the *ger* gene. Unexpectedly, in addition to 1-*O*-MEGs, we were now also able to detect small relative abundances of diether lipids with alkyl chains derived from fatty acids of *E. coli* (Table 2). Hence, the biosynthesis of diether lipids previously hypothesized to involve two independent enzymes, was found to occur solely through the expression of Ger protein, as revealed by our *in vivo* gene-expression results. Further experiments will be required to understand if the conversion of ester to ether bonds indistinctively happens in all phospholipid species or if there are preferred substrates for the Ger enzyme to produce mono and diether lipids. However, we noted already a remarkable difference between PEs and PGs with respect to the presence of the DAG, AEG, and DEG core lipids, with PGs containing relatively more ether bonds in their core (Figure 3B). This likely indicates that PGs and PEs are modified themselves by the activity of Ger and not their precursor, CDP-DAG (Figure 13) and that the degree of conversion of ester into ether bonds depends on phospholipid type. The much lower abundance of 2-*O*-MEGs in comparison to 1-*O*-MEGs in *D. alkenivorans* (Grossi et al., 2015; Vinçon-Laugier et al., 2017) suggests that the transformation of ester into ether bonds by the Ger enzyme starts preferentially at the *sn*-1 position and then proceeds at the *sn*-2 position (Figure 13).

**Figure 13.**
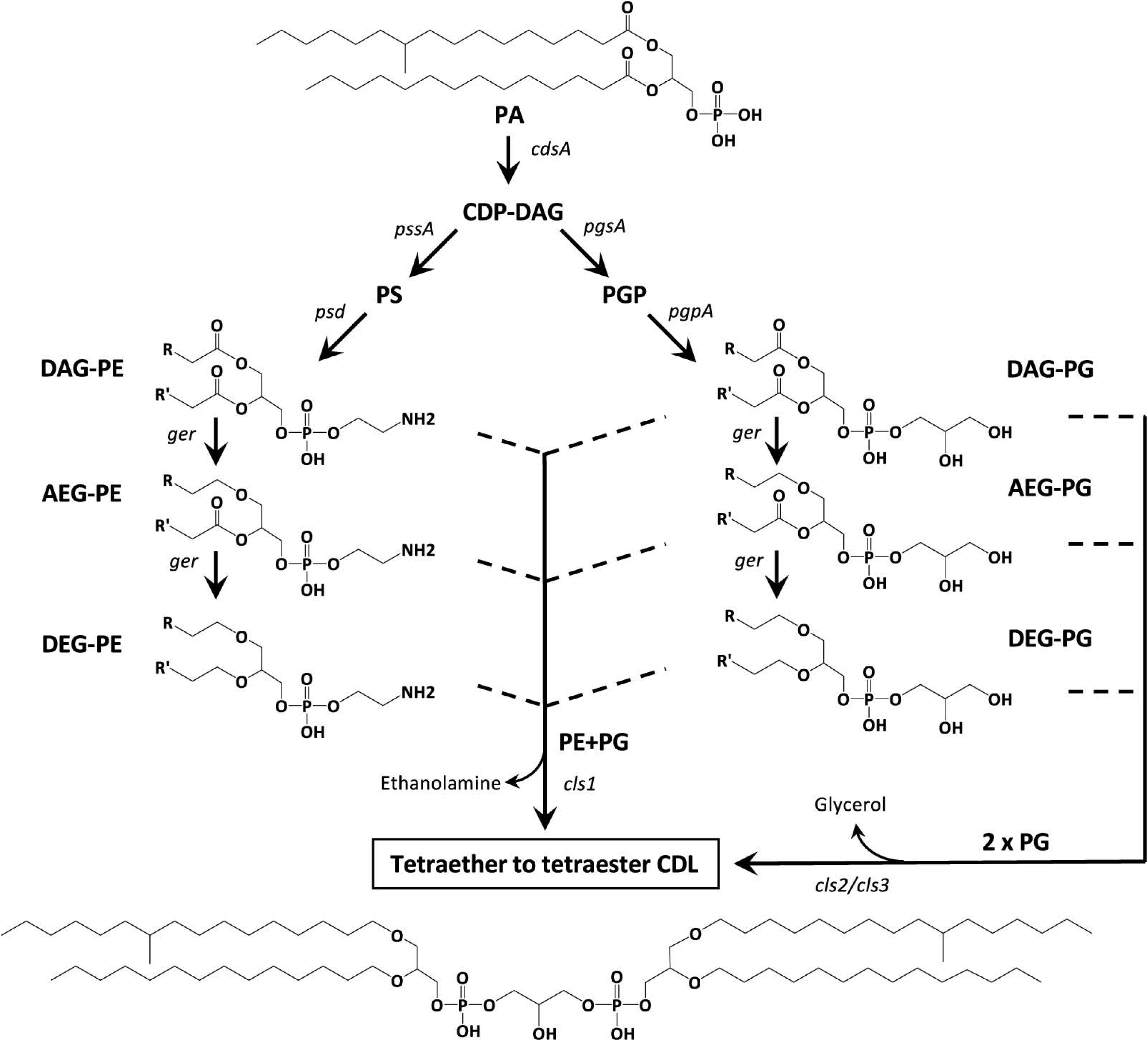
Proposed biosynthetic pathways for the formation of mono- to tetra-alkyl ether CDLs in *D. alkenivorans* based on the lipids and enzymes (Supplementary Table S1) detected in this work. PG and PE are synthesized by the canonical phospholipid biosynthetic pathway with CDP-DAG as the substrate. The diacyl (DAG) core of PG and/or PE can be converted into 1-alkyl, 2-acylglycerol core (AEG) and successively to 1,2-dialkylglycerol (DEG) core by a glycerol ether reductase enzyme (Ger). Cardiolipins can then be produced by a cardiolipin synthase (Cls) by two pathways involving either 2 PGs (as DAG, AEG or DEG) or a PG and a PE (both as DAG, AEG and DEG). The structural genes encoding the enzymes responsible for each step are indicated. Illustrated structures are examples of characteristic lipids formed by *D. alkenivorans* grown on *n*-hexadec-1-ene.

### Occurrence and Potential Formation of Lyso-phospholipids

Our UHPLC-HRMS^2^ revealed a complex mixture of lyso-PEs, PGs and CDLs (Figures 3 and 10). Lyso-lipids are typically thought to be formed by hydrolysis of one of the acyl chains, either by intrinsic phospholipases (Sahonero-Canavesi et al., 2019), or by chemical hydrolysis during the extraction. Phospholipases A/patatin-like are encoded by the genome of *D. alkenivorans* (Table S1). Typically, lysolipids are considered to be toxic for organisms as they may impair the membrane organization, but lysolipids are also considered a ‘signal’ of lipid remodelling (Cao et al., 2004).

The structure of the (di)lyso phospholipids in *D. alkenivorans* reveals that they can readily be explained by the hydrolysis of the ester bonds of the various PEs, PGs, and CDLs. It is noteworthy that the relative abundance of lyso components varies markedly within these three groups of phospholipids (Figure 3): CDLs have the highest abundance (38 % of ‘total CDLs’) and PGs the lowest abundance (9 % of ‘total PGs’) (Figure 3). This is most probably a direct consequence of the percentage of ester/ether bonds in their structures since phospholipids can only lose their acyl groups by hydrolysis to form lyso components. PGs have only 28% ester bonds in their core lipids, which is substantially lower than for PEs (60%). CDLs have an even higher chance to lose an acyl substituent because they are composed of two core lipids.

## Conclusions

The lipidome analysis by UHPLC-HRMS^n^ of the alkylglycerol-containing mesophilic sulfate-reducing bacterium *D. alkenivorans* has led to the identification of a mixture of CDLs containing one to four ether-linked alkyl chains (i.e., from monoether/triester to tetraether), further extending the structural diversity of this complex family of membrane lipids occurring widespread in all three domains of life.

The high proportion of alkyl ether CDLs associated with more conventional alkyl ether PEs and PGs, each accounting for around a third of total phospholipids in this bacterium, suggests an important role/function of these novel lipid structures in the bacterial membrane, which undoubtedly merits further investigation. The remodeling of the *D. alkenivorans* lipidome under various nutrient stresses and physico-chemical perturbations is currently under investigation.

The identification of three CDL synthases in the genome of *D. alkenivorans* resembling those previously reported *in E. coli* and theoretical calculation based on the alkyl ether composition of PEs and PGs support two modes of alkyl/acyl CDL biosynthesis, involving the condensation of two PG molecules or a PG and a PE molecule, as previously proposed in many bacteria. These bacterial biosynthetic pathways, from DEGs, AEGs or DAGs with a PE or PG headgroup to mixed ether/ester CDLs, highlight the promiscuity of prokaryotic CDL synthases.

Finally, our heterologous gene expression experiment has shown that expression of Ger protein alone may be sufficient to reduce the two ester bonds present in DAG-PEs and -PGs successively at the *sn*-1 and the *sn*-2 positions, allowing the first biosynthetic pathways to bacterial alkyl ether CDLs to be proposed. It is likely, however, that a broader set of enzymes may be involved in the production of bacterial (di)ether phospholipids, a point that will certainly needs to be addressed in the future.

## Author contributions for manuscript

**E. C. Hopmans**: Conceptualization, Data curation, Formal analysis, Investigation, Methodology, Validation, Visualization, Writing & Editing. **V. Grossi**: Conceptualization, Formal analysis, Funding acquisition, Investigation, Methodology, Validation, Visualization, Writing & Editing. **D.X. Sahonero-Canavesi**: Formal analysis, Investigation, Methodology, Validation, Writing & Editing. **N. Bale**: Investigation, Methodology, Validation, Editing. **C. Cravo-Laureau**: Formal analysis, Investigation, Editing. **J. S. Sinninghe Damsté**: Conceptualization, Funding acquisition, Investigation, Methodology, Validation, Visualization, Writing & Editing.

## Acknowledgements

We thank I. Mitteau for help with bacterial cultures., M. Koenen for experimental assistance with lipid analysis, D. Dorhout and M. Verweij for analytical support, L. Strack van Schijndel for experimental assistance with expression experiments and Dr. L. Villanueva for helpful discussions.

## Supplementary figures

**Figure S1.** Cardiolipins and lyso-cardiolipins with ether-bound alkyl chains. **A)** tetraether cardiolipin with 62 alkyl carbon atoms composed of two C_31_DEG with C_17_ and C_14_ alkyl chains; **B)** monolyso-cardiolipin with 48 alkyl carbon atoms composed of C_14_/C_17_DEG + C_17_MEG (peak **n**); **C)** monolyso-cardiolipin with 45 alkyl carbon atoms composed of C_14_/C_17_DEG + C_14_MEG; **D)** dilyso-cardiolipin with 34 alkyl carbon atoms composed of two C_17_MEG (C_17_/C_17_DEG was not detected); **E)** dilyso-cardiolipins with 31 alkyl carbon atoms composed of C_14_MEG+C_17_MEG or C_14_/C_17_DEG ; **F)** dilyso-cardiolipins with 28 alkyl carbon atoms composed of two C_14_MEG or C_14_/C_14_DEG (peak **p2**). Diagnostic spectra of **C** and **E** are shown in figure 8.

**Figure S2.** Multiple sequence alignment of the three potential cardiolipin synthases Cls from *D. alkenivorans* and the bacterial-type Cls belonging to the PLD family characterized in bacterial species (*Escherichia coli, Staphylococcus aureus, Enterococcus faecalis, Pseudomonas fluorescens, Bacillus subtilis, Agrobacterium tumefaciens, Helicobacter pylori, Pseudomonas syringae*), archaea (*Methanospirillum hungatei*) and eukarya (*Trypanosoma brucei brucei*). The partial amino acid sequences of the three putative *D. alkenivorans* Cls containing the two conserved HXKXXXXD motifs (black boxes), the conserved glycines (black stars) and other strictly conserved residues N, D, Y, S and E (black circles) as all other Cls sequences. The amino acid consensus is colored according to 50% identity conservation. The consensus sequence logo is shown under both alignments. Four blocks with the conserved sequence are shown.

**Figure S3.** Multiple sequence alignment of the three potential cardiolipin synthases Cls from *D. alkenivorans* and the bacterial-type Cls belonging to the PLD family characterized in bacterial species (*Escherichia coli, Staphylococcus aureus, Enterococcus faecalis, Pseudomonas fluorescens, Bacillus subtilis, Agrobacterium tumefaciens, Helicobacter pylori, Pseudomonas syringae*), archaea (*Methanospirillum hungatei*) and found in eukaryotes (*Trypanosoma brucei brucei*). Complete amino acid sequences showing the conserved regions along all the cardiolipin synthases (characterized and potential) The residues are colored according to 50% identity conservation. The consensus sequence logo is shown under the alignments.

