## supplemental figures and tables for "Mono- to tetra-alkyl ether cardiolipins in a mesophilic, sulfate-reducing bacterium identified by UHPLC-HRMS^n^: A novel class of membrane lipids"

### Supplementary Figures and Tables

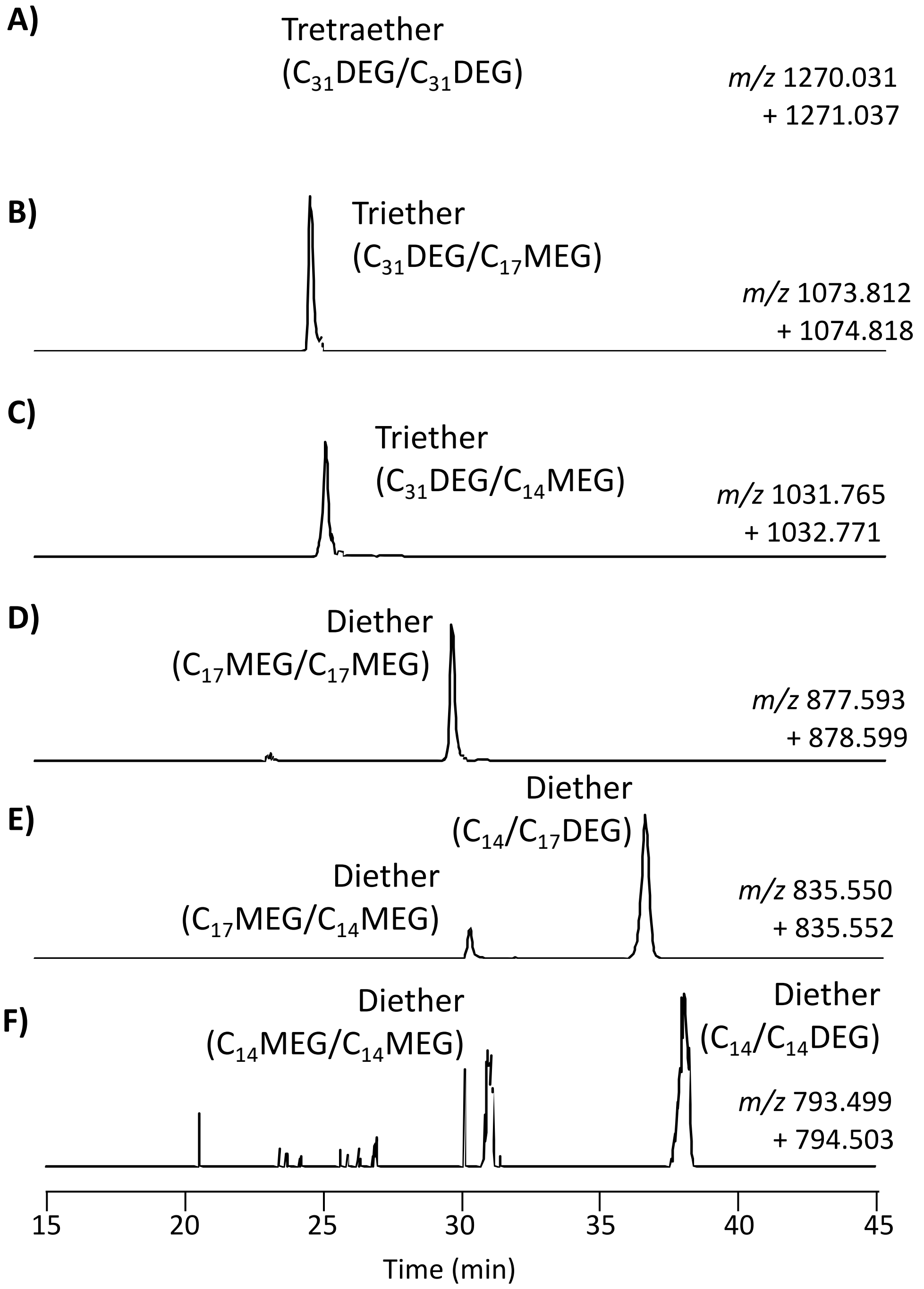


**Figure S1.** Cardiolipins and lyso-cardiolipins with ether-bound alkyl chains. **A)** tetraether cardiolipin with 62 alkyl carbon atoms composed of two $C_{31}$DEG with $C_{17}$ and $C_{14}$ alkyl chains; **B)** monolyso-cardiolipin with 48 alkyl carbon atoms composed of $C_{14}$/$C_{17}$DEG + $C_{17}$MEG (peak **n**); **C)** monolyso-cardiolipin with 45 alkyl carbon atoms composed of $C_{14}$/$C_{17}$DEG + $C_{14}$MEG; **D)** dilyso-cardiolipin with 34 alkyl carbon atoms composed of two $C_{17}$MEG ($C_{17}$/$C_{17}$DEG was not detected); **E)** dilyso-cardiolipins with 31 alkyl carbon atoms composed of $C_{14}$MEG+$C_{17}$MEG or $C_{14}$/$C_{17}$DEG ; **F)** dilyso-cardiolipins with 28 alkyl carbon atoms composed of two $C_{14}$MEG or $C_{14}$/$C_{14}$DEG (peak **p2**). Diagnostic spectra of **C** and **E** are shown in figure 8.

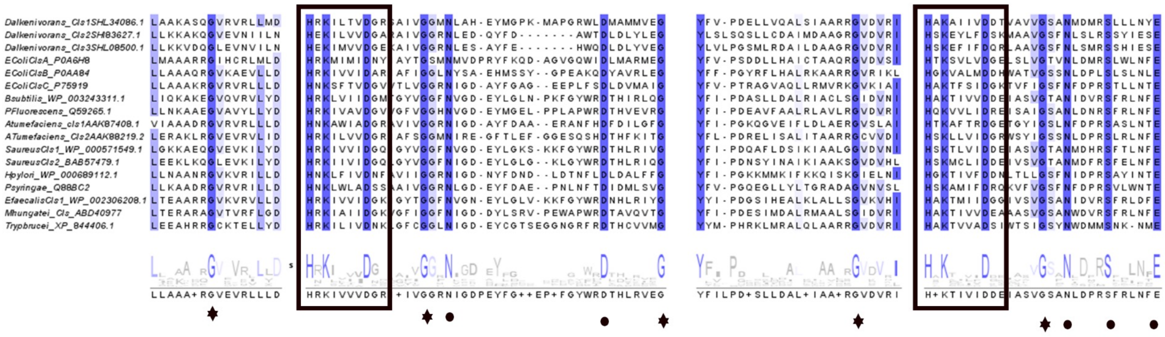

**Figure S2.** Multiple sequence alignment of the three potential cardiolipin synthases Cls from *D. alkenivorans* and the bacterial-type Cls belonging to the PLD family characterized in bacterial species (*Escherichia coli, Staphylococcus aureus, Enterococcus faecalis, Pseudomonas fluorescens, Bacillus subtilis, Agrobacterium tumefaciens, Helicobacter pylori, Pseudomonas syringae*), archaea (*Methanospirillum hungatei*) and eukarya (*Trypanosoma brucei brucei*). The partial amino acid sequences of the three putative *D. alkenivorans* Cls containing the two conserved HXKXXXXD motifs (black boxes), the conserved glycines (black stars) and other strictly conserved residues N, D, Y, S and E (black circles) as all other Cls sequences. The amino acid consensus is colored according to 50% identity conservation. The consensus sequence logo is shown under both alignments. Four blocks with the conserved sequence are shown.

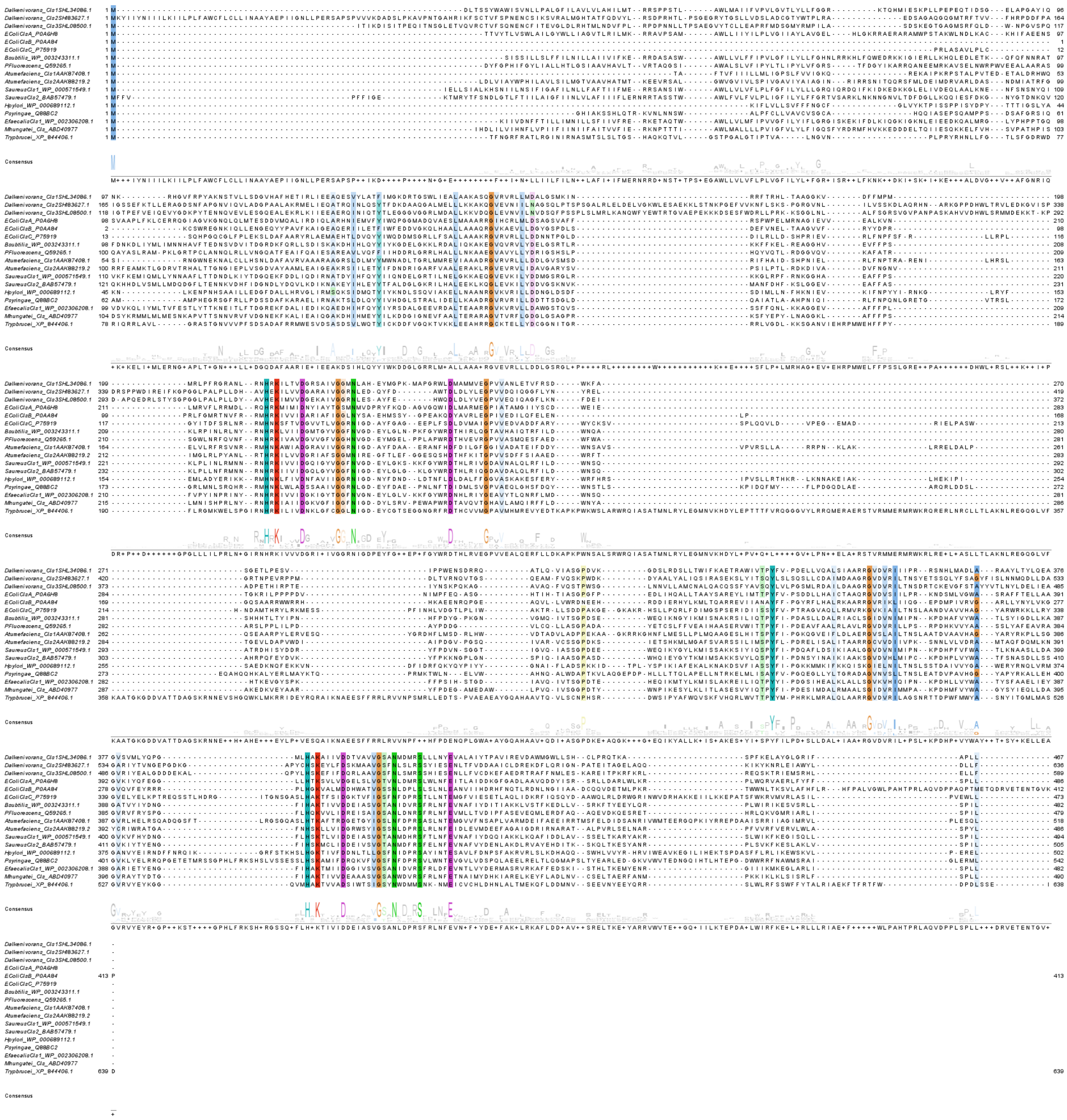

**Figure S3.** Multiple sequence alignment of the three potential cardiolipin synthases Cls from *D. alkenivorans* and the bacterial-type Cls belonging to the PLD family characterized in bacterial species (*Escherichia coli, Staphylococcus aureus, Enterococcus faecalis, Pseudomonas fluorescens, Bacillus subtilis, Agrobacterium tumefaciens, Helicobacter pylori, Pseudomonas syringae*), archaea (*Methanospirillum hungatei*) and found in eukaryotes (*Trypanosoma brucei brucei*). Complete amino acid sequences showing the conserved regions along all the cardiolipin synthases (characterized and potential) The residues are colored according to 50% identity conservation. The consensus sequence logo is shown under the alignments.

**Table S1.** Potential genes and associated enzymes involved in the synthesis of bacterial alkyl ether CDL and their lyso-counterparts encoded by the genome of *D. alkenivorans*.

| Gene | Enzyme Description | Uniprot accession | NCBI accesion (Ref. Seq.) | Number of transmem-brane domains[a] | Membrane protein |
|---|---|---|---|---|---|
| *cls1* | Cardiolipin synthase 1 | SHL34086.1 | WP_073479011.1 | 2 | Yes |
| *cls2* | Cardiolipin synthase 2 | SHI83627.1 | WP_139264592.1 | 1 | Yes |
| *cls3* | Cardiolipin synthase 3 | SHL08500.1 | WP_073478579.1 | ND | Yes |
| *ymdB* | O-acetyl-ADP-ribose deacetylase | SHL16402.1 | WP_073478717.1 | | |
| *ger* | Glycerol ester reductase | SHJ90043.1 | WP_073476268.1 | 1* | Yes |
| n.d. | Potential Glycerol ester reductase 2 | SHK01260.1 | WP_073476658.1 | ND | Yes |
| *cdsA* | phosphatidate cytidylyltransferase | SHI78129.1 | WP_073472543.1 | 6 | Yes |
| *pgsA* | CDP-diacylglycerol-glycerol-3-phosphate 3-phosphatidyltransferase | SHK52233.1 | WP_073477644.1 | 5 | Yes |
| *pgpA* | Phosphatidylglycerophosphatase A | SHJ07628.1 | WP_073473540.1 | 3 | Yes |
| *pssA* | CDP-diacylglycerol-serine O-phosphatidyltransferase | SHL43585.1 | WP_083611310.1 | 6 | Yes |
| *psd* | phosphatidylserine decarboxylase family protein | SHL43626.1 | WP_073479141.1 | 2 | Yes |
| *Pla1* | phospholipase A1 | SHK83058.1 | WP_073478171.1 | 1 | Yes |
| | Patatin-like phospholipase 1 | SHJ82754.1 | WP_073475982.1 | | |

[a] transmembrane domains were detected by application of the TMHMM algorithm (Krogh et al., 2001)

**Table S2.** The sequence similarity as determined by pBLASTof ClsA, ClsB, ClsC, archaeal Cls compared with proteins encoded by the genomes of *D. alkenivorans* and the closely related species *D. aliphaticivorans*.

| Bacterium | | | *E. coli* | | *M. hungatei* | *S. coelicolor* | *D. alkenivorans* |
|---|---|---|---|---|---|---|---|
| Protein | | **ClsA** P0A6H8 (486 AA) | **ClsB** P0AA84 (413 AA) | **ClsC** P75919 (473 AA) | **arCls** A0A8F5VN54 (504 AA) | Sco1389 (215 AA) | **Ger** SHJ90043 (1458 AA) |
| *Desulfatibacillum alkenivorans* | | | | | | | |
| WP_073479011.1 | Evalue | 0 | 2E-143 | 1E-127 | 0 | | |
| (467 AA) | query cover (%) | 88 | 76 | 82 | 93 | | |
| | similarity (%) | 30.9 | 32.5 | 25.6 | 30.2 | | |
| WP_139264592.1 | Evalue | 2E-112 | 3E-74 | 6E-86 | 6E-105 | | |
| (612 AA) | query cover (%) | 72 | 73 | 65 | 75 | | |
| | similarity (%) | 25.9 | 20.9 | 21.2 | 26.2 | | |
| WP_073478579.1 | Evalue | 9E-86 | 9E-85 | 3E-81 | 1E-117 | | |
| (589 AA) | query cover (%) | 66 | 53 | 63 | 57 | | |
| | similarity (%) | 27.3 | 24.7 | 23.5 | 25.0 | | |
| WP_073477644.1 | Evalue | | | | | 6E-56 | |
| (195 AA) | query cover (%) | | | | | 56 | |
| | similarity (%) | | | | | 34.4 | |
| SHK01260 | Evalue | | | | | | 0 |
| (1458 AA) | query cover (%) | | | | | | 96 |
| | similarity (%) | | | | | | 25.9 |
| *Desulfatibacillum aliphaticivorans* | | | | | | | |
| WP_015948190.1 | Evalue | 0 | 1E-141 | 4E-125 | 0 | | |
| (467 AA) | query cover (%) | 88 | 76 | 82 | 93 | | |
| | similarity (%) | 31.3 | 32.5 | 25.1 | 30.4 | | |
| WP_136360700.1 | Evalue | 8E-114 | 5E-76 | 1E-84 | 3E-106 | | |
| (612 AA) | query cover (%) | 74 | 73 | 65 | 75 | | |
| | similarity (%) | 25.9 | 21.3 | 20.6 | 26.8 | | |
| WP_012610398.1 | Evalue | 3E-84 | 9E-85 | 5E-83 | 3E-116 | | |
| (589 AA) | query cover (%) | 67 | 53 | 63 | 71 | | |
| | similarity (%) | 27.0 | 24.7 | 23.5 | 25.3 | | |
| WP_012609567.1 | Evalue | | | | | 4E-56 | |
| (195 AA) | query cover (%) | | | | | 56 | |
| | similarity (%) | | | | | 35.3 | |
| WP_028315991.1 | Evalue | | | | | | 0 |
| (1401 AA) | query cover (%) | | | | | | 96 |
| | similarity (%) | | | | | | 25.2 |
